# Polarized vesicle transport requires AP-1-mediated recruitment of KIF13A and KIF13B at the trans-Golgi network

**DOI:** 10.1101/2023.04.06.535716

**Authors:** Andrew C. Montgomery, Christina S. Mendoza, Alex Garbouchian, Geraldine B. Quinones, Marvin Bentley

## Abstract

Neurons are polarized cells that require accurate membrane trafficking to maintain distinct protein complements at dendritic and axonal membranes. The Kinesin-3 family members KIF13A and KIF13B are thought to mediate dendrite-selective transport, but the mechanism by which they are recruited to polarized vesicles and the differences in the specific trafficking role of each KIF13 have not been defined. We performed live-cell imaging in cultured hippocampal neurons and found that KIF13A is a dedicated dendrite-selective kinesin. KIF13B confers two different transport modes, both dendrite- and axon-selective transport. Both KIF13s are maintained at the trans-Golgi network by interactions with the heterotetrameric adaptor protein complex AP-1. Interference with KIF13 binding to AP-1 resulted in disruptions to both dendrite- and axon-selective trafficking. We conclude that AP-1 is the molecular link between the sorting of polarized cargoes into vesicles and the recruitment of kinesins that confer polarized transport.

## INTRODUCTION

The neuron is the fundamental unit of the nervous system that mediates electrochemical signaling. Neurons are polarized cells that typically have multiple dendrites and one axon (Bentley and Banker, 2016; Craig and Banker, 1994). Dendrites and axons perform different functions in neuronal signaling and consequently require distinctive complements of membrane proteins. Polarized membrane proteins are delivered to their proper domain by membrane trafficking. Despite their fundamental importance, the trafficking mechanisms that maintain neuronal polarity are still unclear (Bentley and Banker, 2016; Radler et al., 2020).

Polarized transmembrane proteins are moved in vesicles that undergo selective transport (Burack et al., 2000): dendrite-selective vesicles move bidirectionally in dendrites and are restricted from entering the axon (Burack et al., 2000; Silverman et al., 2001; Al-Bassam et al., 2012; Jensen et al., 2014; Petersen et al., 2014; Frank et al., 2020; Eichel et al., 2022); axon-selective vesicles can move bidirectionally in dendrites, but preferentially enter the axon where they undergo mostly anterograde movement (Burack et al., 2000; Petersen et al., 2014; Nabb and Bentley, 2022). Polarized transmembrane proteins are synthesized in the endoplasmic reticulum and modified in the Golgi apparatus. At the trans-Golgi network (TGN), proteins are sorted into vesicles that deliver them to their destination (Bonifacino, 2014; Guardia et al., 2018; Ford et al., 2021; Ramazanov et al., 2021). Newly synthesized dendritically polarized proteins are sorted by the heterotetrameric clathrin adaptor protein complex 1 (AP-1) (Farías et al., 2012). In addition to Golgi-derived vesicles, a subpopulation of endocytic vesicles undergoes dendrite-selective transport (Frank et al., 2020). In contrast, the machinery that mediates axonal sorting has not been defined in mammalian neurons.

Dendrite-selective vesicles are transported by two motors of the Kinesin-3 family, KIF13A and KIF13B (Jenkins et al., 2012). In neurons, KIF13A binds dendritically polarized vesicles (Yang et al., 2019; Frank et al., 2020) and mediates localization of AMPA (Gutiérrez et al., 2021) and serotonin (Zhou et al., 2013) receptors in dendrites. In non-neuronal cells, KIF13A participates in the transport and positioning of endosomes (Nakagawa et al., 2000; Delevoye et al., 2009, 2014; Bentley et al., 2015; Etoh and Fukuda, 2019). KIF13B similarly binds endosomes in non-neuronal cells (Bentley et al., 2015; Mills et al., 2019). In addition to its role in dendrite-selective transport (Frank et al., 2020), KIF13B is also required for axon selection during neuronal development (Horiguchi et al., 2006; Yoshimura et al., 2010; Yu et al., 2020). Despite this progress, the specific functions of KIF13A and KIF13B in polarized vesicle transport—and the specific vesicle subpopulations they move—have not been defined.

A vesicle transport event is initiated by two steps: 1) cargo sorting and vesicle budding from the donor compartment, and 2) motor-based transport to the cellular destination. Because these steps are strictly successive, they must be mechanistically linked; that is, the proteins required for proper transport must be determined during vesicle formation. For example, a vesicle containing proteins destined for the dendritic plasma membrane must be *programmed* to recruit the correct complement of molecular motors and motor regulatory proteins that confer dendrite-selective transport. Despite the crucial importance of motors to vesicle transport, little is known about how appropriate kinesins are recruited to specific vesicle populations and how adaptors regulate vesicle-bound kinesins (Nabb et al., 2020).

We performed live-cell imaging in cultured hippocampal neurons to quantitatively characterize the functions of KIF13A and KIF13B in polarized vesicle transport. We found that KIF13A is dedicated to transporting both Golgi-derived and endocytosed dendrite-selective vesicles. KIF13B participates in the transport of Golgi-derived and endocytosed dendrite-selective vesicles but performs a second function in mediating transport of axon-selective vesicles. These data show that KIF13B transports different vesicles with distinct transport behaviors, providing strong evidence for on-vesicle regulation (Nabb et al., 2020). Furthermore, neurons maintain pools of KIF13A and KIF13B at the TGN by binding to AP-1 and disrupting these interactions diminishes both dendrite- and axon-selective trafficking. This suggests that AP-1 performs cargo sorting into polarized vesicles and mediates recruitment of the proper kinesins. Together, these data show that AP-1 is a molecular link between the sorting of polarized membrane proteins and the recruitment of KIF13s to mediate long-range selective transport.

## RESULTS

### KIF13A specializes in dendrite-selective transport

Two Kinesin-3 family members, KIF13A and KIF13B, mediate dendrite-selective transport in neurons (Jenkins et al., 2012; Yang et al., 2019; Gutiérrez et al., 2021), but the extent to which they are complementary or redundant is unclear. Vesicles containing important dendritic membrane proteins such as mGluR1, AMPA receptors, and NMDA receptors can be conveniently labeled by expressing fluorescently-tagged transferrin receptor (TfR) (Burack et al., 2000; Silverman et al., 2001; Wang et al., 2008; Farías et al., 2012, 2015; Frank et al., 2020). However, the pool of “dendrite-selective vesicles”—even those visualized with a single fluorescent marker— consist of distinct vesicle subpopulations. TfR traffics in at least two different subpopulations: 1) Golgi-derived vesicles that contain newly synthesized TfR that transport from the Golgi to the plasma membrane and 2) endocytic vesicles that contain TfR that cycles between the plasma membrane and dendrite-selective endosomes (de Jong et al., 1990; Harding et al., 1983). While we previously found that KIF13A and KIF13B bind TfR vesicles, those experiments could not distinguish between subpopulations (Jenkins et al., 2012; Bentley and Banker, 2015). Furthermore, the relative amounts of Golgi-derived and endocytic TfR vesicles are not known. Therefore, it is unclear if KIF13A and KIF13B are functionally redundant or specialize in specific aspects of dendrite-selective trafficking.

To determine if KIF13A and KIF13B specialize in different aspects of dendrite-selective transport, we used a technique we developed for visualizing vesicle-bound kinesins in cultured hippocampal neurons (Yang et al., 2019; Frank et al., 2020; Montgomery et al., 2022; Garbouchian et al., 2022). We coexpressed Halo-KIF13A or Halo-KIF13B tail with TfR-GFP and a SignalSequence (SigSeq)-mCherry construct (Fig. 1). SigSeq-mCherry consists of the signal sequence from neuropeptide-Y fused to mCherry which targets the fluorophore to the lumen of Golgi-derived vesicles (El Meskini et al., 2001; Kaech et al., 2012a; Das et al., 2013; Fang et al., 2014; Ganguly et al., 2015, 2017; Nabb and Bentley, 2022). When Golgi-derived mCherry-containing vesicles fuse with the plasma membrane, the fluorophore is released into the culture medium. With this approach, TfR-GFP vesicles colabeled with mCherry are Golgi-derived, whereas TfR-GFP vesicles without mCherry are endocytic (Fig. 1A).

**Figure 1:**
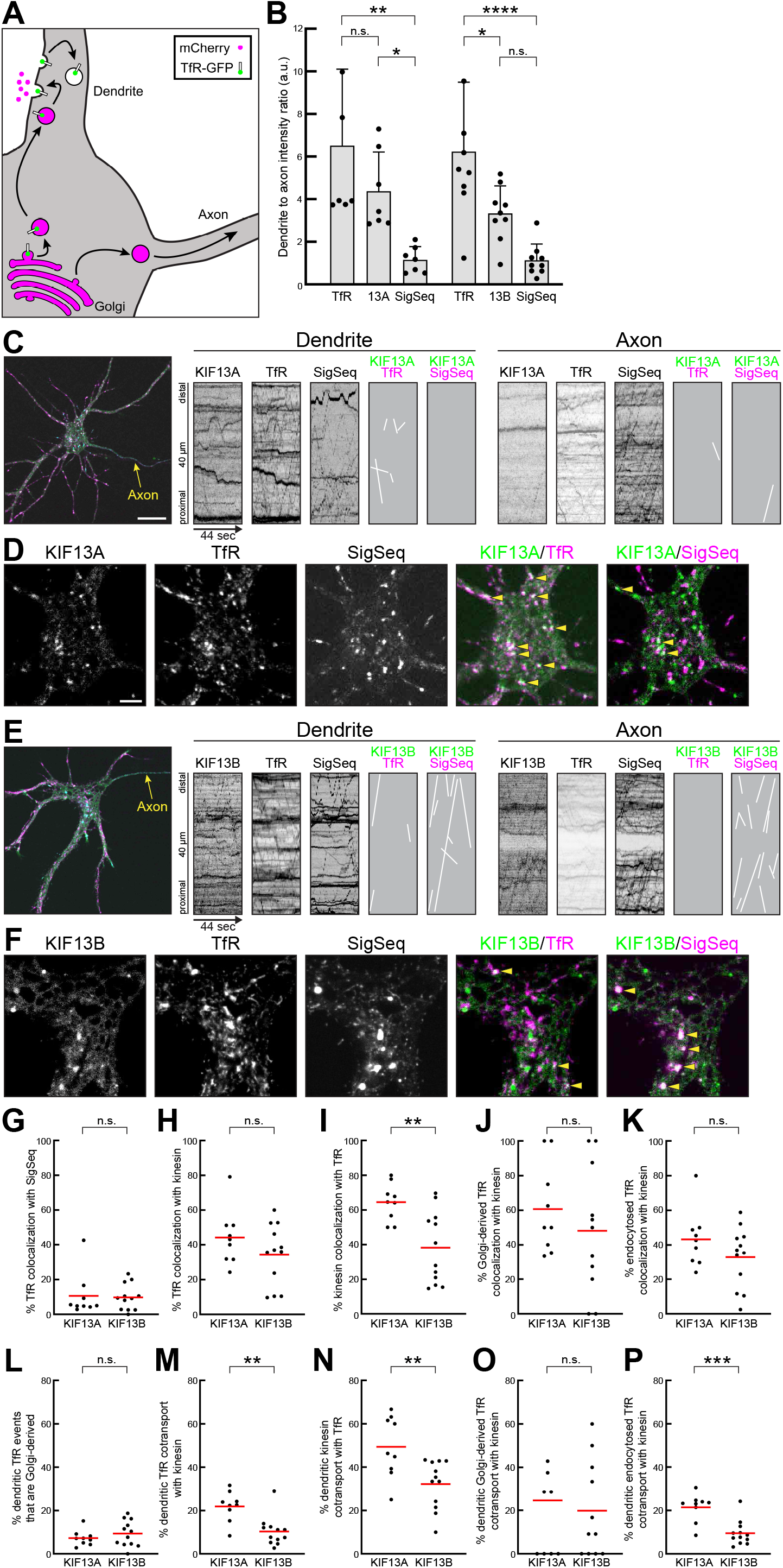
Closely related kinesins KIF13A and KIF13B exhibit different neuronal localization and transport. (A) Schematic describing the labeling strategy to visualize Golgi-derived vesicles. Expression of SigSeq-mCherry results in visualization of the lumen of the secretory pathway by mCherry. Upon vesicle fusion with the plasma membrane, mCherry is released into the culture medium. Endocytic vesicles are not labeled by this strategy. (B) Polarity measurements for indicated proteins. Each TfR condition has one datapoint omitted that exceeded the y-axis. Cell numbers: KIF13A: 7; KIF13B: 9. (C-F) Representative images and kymographs of 9 DIV hippocampal neurons expressing either Halo-KIF13A tail (C&D) or Halo-KIF13B tail (E&F) with SiqSeq-mCherry to visualize Golgi-derived vesicles, and transferrin receptor-GFP. Halotags were visualized with JF646. Kymographs show transport from a dendrite and the axon and cotransport events were redrawn for clarity. High magnification images show the soma of each neuron. Arrowheads indicate examples of colocalization. Low magnification scale bar: 20 µm. High magnification soma scale bar: 5 µm. (G-K) Quantification of colocalization between indicated constructs in the somata. (L-P) Kymograph cotransport analysis for the indicated constructs in the dendrites. One datapoint was omitted in (O) from the KIF13A condition that exceeded the y-axis. Sample size: KIF13A: 9 cells; KIF13B: 12 cells. *p<0.05; **p<0.01; ***p<0.001.

We first measured protein polarity of each construct (Fig. 1B). As expected, TfR and KIF13A were polarized to dendrites. KIF13B was less polarized and SigSeq was unpolarized, which is consistent with it labeling Golgi-derived vesicles that travel to all neurites. We generated kymographs to analyze vesicle transport (Fig. 1C&E). Kymograph lines with a positive slope indicate anterograde transport and lines with a negative slope indicate retrograde transport, a convention that is followed in all figures. For clarity, cotransport events from kymographs were redrawn as solid white lines. The movement of each vesicle population reflected its overall polarity (Fig. 1C&E). TfR and KIF13A vesicles moved in dendrites but were largely absent from axons. SigSeq underwent bidirectional movements in dendrites and exhibited mostly anterograde movement in the axon. KIF13B vesicles moved bidirectionally in dendrites, consistent with KIF13B’s function in dendrite-selective transport. Notably, there was also an axonal population of KIF13B-labeled structures, which is consistent with the hypothesis that KIF13B performs an axonal role beyond dendrite-selective vesicle transport.

We evaluated colocalization in somata (Fig. 1D&F), which are biochemically indistinct from dendrites (Moore and Baleja, 2012; Leterrier and Dargent, 2014; Craig and Banker, 1994). Because this analysis was aimed at transport vesicles, we chose imaging planes close to the coverslip that typically contain no Golgi structures. KIF13A colocalized substantially with TfR, but relatively little with SigSeq (Fig. 1D). In contrast, KIF13B colocalized less with TfR than KIF13A did, but more with SigSeq (Fig. 1F).

We next performed a series of quantitative colocalization and cotransport analyses (Fig. 1G-P). Our approach allowed us to measure relative amounts of Golgi-derived (SigSeq positive) and endocytic (SigSeq negative) TfR subpopulations for the first time. Colocalization quantification shows that ∼10 % of TfR was in Golgi-derived vesicles (Fig. 1G), indicating that most TfR vesicles are part of the endocytic recycling pathway. We next measured the fraction of total TfR vesicles that colocalized with each KIF13 (Fig. 1H). 44 % colocalized with KIF13A and 34 % with KIF13B (Fig. 1H). While this difference was not statistically significant, it is part of an emerging trend of KIF13A consistently colocalizing and cotransporting more with TfR than KIF13B. We then measured the fraction of KIF13 vesicles that colocalized with TfR (Fig. 1I). Most KIF13A vesicles (64 %) colocalized with TfR, indicating that dendrite-selective vesicles are the primary KIF13A cargo. In contrast, only 38 % of KIF13B vesicles colocalized with TfR. These data show that KIF13A specializes in dendrite-selective transport. While KIF13B plays some role in dendrite-selective transport, that accounts for significantly fewer of its transport duties compared to KIF13A.

We then measured colocalization of KIF13A or KIF13B with Golgi-derived (identified by TfR-GFP and SigSeq labeling) or endocytic (identified by TfR-GFP labeling without SigSeq) TfR vesicles. Both kinesins colocalized with Golgi-derived (KIF13A: 61 %; KIF13B: 48 %) (Fig. 1J) and endocytosed TfR (KIF13A: 43 %; KIF13B: 33 %) (Fig. 1K). KIF13A consistently displayed higher colocalization with TfR. While the differences between KIF13A and KIF13B were not statistically significant, they support the model that KIF13A is specialized for dendrite-selective transport. In contrast, dendrite-selective organelles consistently accounted for a smaller percentage of KIF13B vesicles.

While colocalization data are instructive, cotransport of two fluorescently tagged proteins is maybe the strongest evidence of two proteins associating with the same vesicle. Furthermore, vesicle transport is a dynamic process and analysis of moving vesicles directly evaluates transport behaviors. We measured the cotransport of different organelles in dendrites (Fig. 1L-P). Results of the cotransport analysis were generally consistent with the colocalization data (Fig. 1G-K). Only ∼8 % of moving TfR vesicles were Golgi-derived (Fig. 1L), closely matching the colocalization analysis (Fig. 1G). From all TfR vesicles, 22 % cotransported with KIF13A and 10 % with KIF13B (Fig. 1M). About half of KIF13A events cotransported with TfR, contrasting KIF13B, which consistently exhibited less cotransport with all TfR populations (Fig. 1N-P).

Visualizing vesicle-associated kinesins in live cells is technically challenging under ideal conditions. Expression of only the vesicle binding tail domain results in a temporal window in which kinesin-labeled vesicles become visible (Yang et al., 2019; Montgomery et al., 2022). The length of that window varies from cell to cell. Achieving vesicle labeling is easiest when only expressing a single kinesin tail. Even then, visible vesicles are probably only a fraction of all organelles bound by specific kinesins. Kinesins are peripheral membrane proteins that cycle on and off vesicles. Therefore, fluorescently tagged kinesins must be expressed at low levels, while in competition with highly expressed endogenous kinesins. Balancing additional expression constructs enhances the difficulty of the experiments. Therefore, the colocalization and cotransport numbers we measured are almost certainly an underestimation of true colocalization and must be interpreted accordingly. Even single digit colocalization and cotransport percentages are likely indicators of substantial biological interaction between a kinesin and a specific vesicle. We sought a complementary approach to determine KIF13A and KIF13B interactions with dendrite-selective endocytic vesicles (Fig. 2A). Neurons expressing either Halo-KIF13A or Halo-KIF13B tail were treated with transferrin (Tf) conjugated to an organic dye. Fluorescent transferrin is endocytosed by endogenous TfR and specifically labels dendrite-selective vesicles in the endocytic pathway (Frank et al., 2020). Tf vesicles were polarized to dendrites (Fig. 2B) where they cotransported with both KIF13A and KIF13B (Fig. 2C&D). Quantification of colocalization and cotransport in dendrites showed that both KIF13A and KIF13B bound Tf vesicles (Fig. 2E-H). A higher proportion of Tf vesicles colocalized with KIF13A (21 %) than KIF13B (12 %) (Fig. 2E), although this was not statistically significant. However, Tf-labeled organelles accounted for a significantly higher fraction of KIF13A vesicles (47 %) than for KIF13B (18 %). We next performed cotransport analyses of Tf with KIF13A or KIF13B. Comparable amounts of Tf vesicles cotransported with KIF13A (11 %) and KIF13B (13 %) (Fig. 2G). Finally, we measured the fraction of KIF13A or KIF13B vesicles that cotransported with Tf and found a significant difference between the two kinesins. KIF13A exhibited significantly more with Tf (29 %) than KIF13B (14 %) (Fig. 2H), which was consistent with the colocalization data (Fig. 2F). These results further highlight the specialization of KIF13A for dendrite-selective transport and show that KIF13B performs additional transport functions.

**Figure 2:**
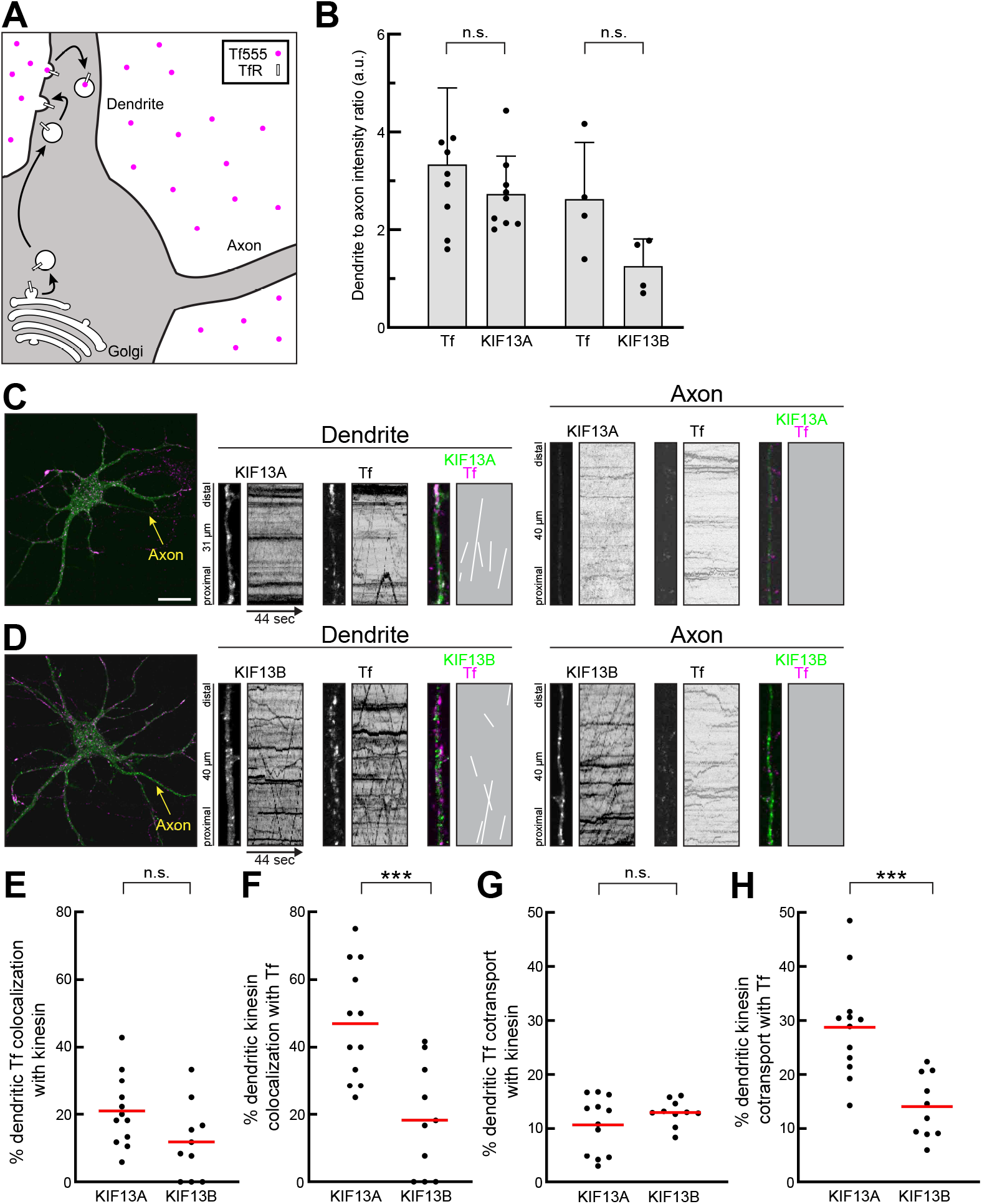
KIF13A preferentially mediates dendrite-selective transport of endocytic vesicles. (A) Schematic describing the labeling strategy to visualize endocytosed vesicles. Fluorescent transferrin (Tf555) in the culture medium binds endogenous transferrin receptor and undergoes endocytosis. Golgi-derived organelles are not labeled by this strategy. (B) Polarity measurements for indicated proteins. One Tf data point was omitted from the KIF13A condition because it exceeded the y-axis. Cell numbers: KIF13A: 9; KIF13B: 4. (C&D) Representative images and kymographs of 8 DIV hippocampal neurons expressing Halo-KIF13A tail (C) or Halo-KIF13B tail (D) and incubated with Tf555. Halotags were visualized with JF646. Kymographs show transport from a dendrite and the axon and cotransport events were redrawn for clarity. Scale bar: 20 µm. (E-H) Quantification of colocalization and cotransport between kinesins and Tf organelles. Sample size: KIF13A: 12 cells; KIF13B: 10 cells. ***p<0.001.

The systematic analyses of dendrite-selective vesicles point to a model where both KIF13A and KIF13B mediate dendrite-selective transport but exhibit different specializations. Our data are consistent with KIF13A being dedicated to dendrite-selective transport. In contrast, we found that KIF13B participates in dendrite-selective transport, although this only accounts for a subset of its vesicles. There is an additional component of KIF13B vesicles—including in axons— that is not dendrite-selective. Therefore, KIF13B may confer different transport parameters that depend on its cargo.

### KIF13B mediates axon-selective transport

KIF13B is thought to participate in axon selection during neuronal development (Yamada et al., 2007; Yoshimura et al., 2010), but the molecular identity of axonal KIF13B vesicles has not been defined. KIF13B vesicles underwent long-range anterograde transport in axons, moved bidirectionally in dendrites, and cotransported with SiqSeq (Fig. 1E). These transport parameters are similar to those we had previously observed of axon-selective vesicles containing neuron-glia cell adhesion molecule (NgCAM) (Nabb and Bentley, 2022; Frank et al., 2022), which led us to hypothesize that KIF13B mediates axon-selective transport. To test this, we coexpressed NgCAM-GFP with Halo-KIF13B tail and performed live-cell imaging (Fig. 3A). Cotransport in dendrites was relatively low, presumably because most dendritic KIF13B vesicles in dendrites are dendrite selective. In contrast, there was anterograde cotransport in axons. Quantification of transport parameters found that KIF13B and NgCAM exhibited similar transport behaviors: vesicles underwent bidirectional movements in dendrites and had a strong anterograde bias in axons (Fig. 3B-E). Overall, we detected fewer KIF13B vesicles than NgCAM vesicles. This is likely because NgCAM is a transmembrane protein that readily labels vesicles whereas KIF13B is a peripheral membrane protein that cycles between vesicle membranes and the cytoplasm.

**Figure 3:**
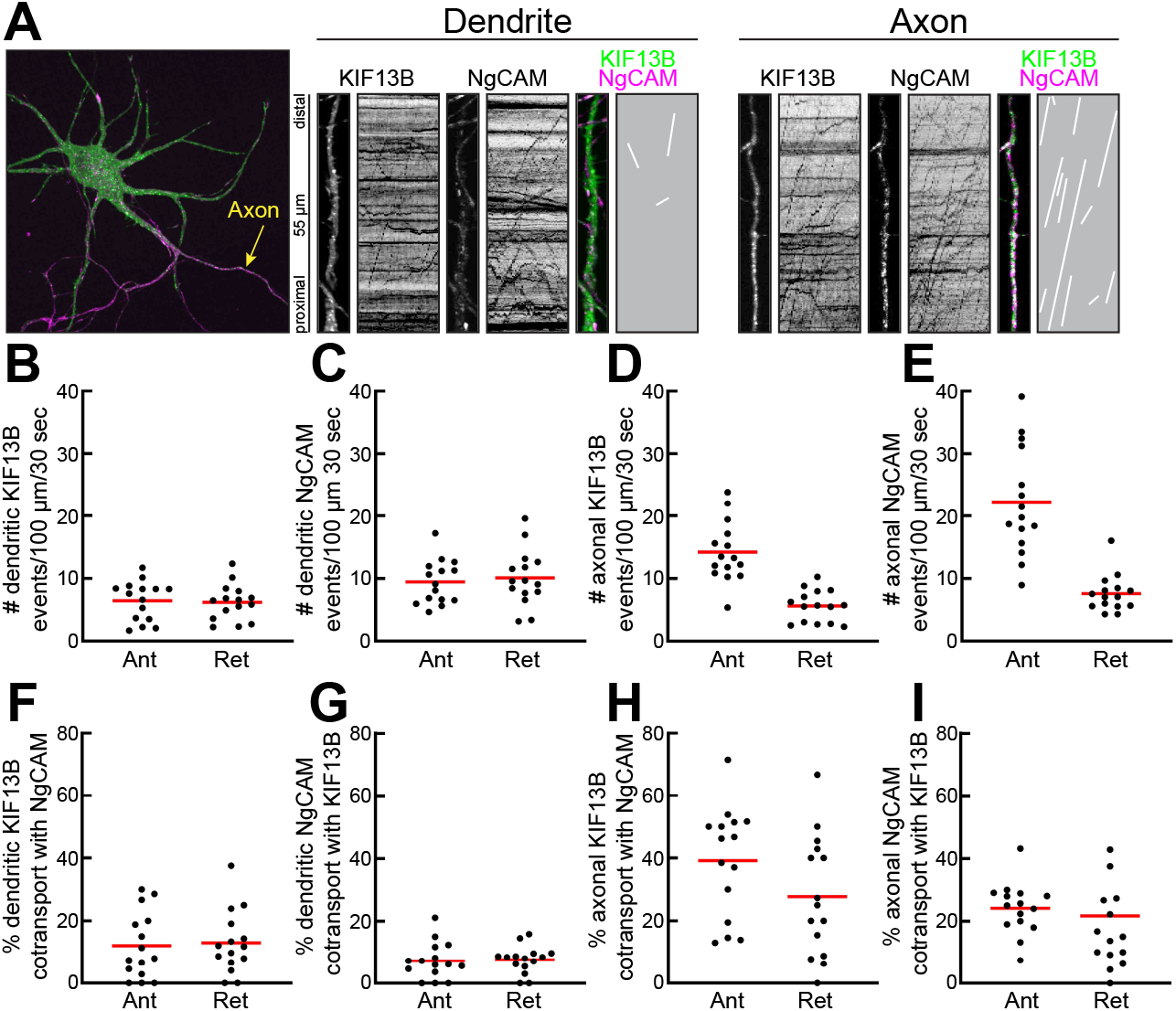
KIF13B transports axon-selective vesicles. (A) Representative image of an 8 DIV hippocampal neuron expressing Halo-KIF13B tail and NgCAM-GFP. Halotags were visualized with JF549. Kymographs show transport from the axon and a dendrite and cotransport events were redrawn for clarity. Scale bar: 20 µm. (B-I) Quantification of transport events and cotransport in the axon and dendrites. One retrograde data point in (I) was omitted because it exceeded the y-axis. Sample size: 15 cells.

We next quantified the cotransport of KIF13B and NgCAM. NgCAM trafficking in dendrites is complex, as there are at least two populations: 1) Golgi-derived NgCAM vesicles that traverse dendrites before reaching the axon and 2) endocytic vesicles containing wayward NgCAM that is being trafficked to lysosomes for degradation (Nabb and Bentley, 2022). Cotransport of KIF13B and NgCAM in dendrites was below 15 % (Fig. 3F&G). In axons nearly all anterograde NgCAM transport is in Golgi-derived vesicles that contain newly-synthesized proteins (Nabb and Bentley, 2022; Frank et al., 2022). We performed the same analyses in axons and found significant cotransport. Here, 39 % of anterograde KIF13B vesicles cotransported with NgCAM and 24 % of anterograde NgCAM vesicles cotransported with KIF13B (Fig. 3H&I). These results show that KIF13B is involved in axonal transport of axon-selective vesicles. It is notable that the assumption in the field has been that Kinesin-1 family members (KIF5s) mediate axon-selective transport (Nabb et al., 2020). However, this assumption was largely due to observations that KIF5 motor domains are axon-selective (Nakata and Hirokawa, 2003; Jacobson et al., 2006; Nakata et al., 2011; Huang and Banker, 2012) and not by detecting kinesin–cargo cotransport. While KIF5 clearly mediates axonal transport of multiple organelles (Wang and Schwarz, 2009; Encalada et al., 2011; Fu and Holzbaur, 2013; Farías et al., 2017; Fukuda et al., 2021), it is possible that KIF13B plays at least a complementary—if not leading—role in the transport of axon-selective vesicles.

### Neurons maintain a stable pool of KIF13B at the trans-Golgi network

Polarized cargoes are sorted into vesicles for selective transport at the trans-Golgi network (TGN) (Farías et al., 2012; Petersen et al., 2014; Nabb and Bentley, 2022). During or after budding, vesicles must acquire the proper kinesin motors. Current understanding of any kinesin’s recruitment is at best fragmentary, and even less is known about the process by which kinesins are recruited to polarized vesicles (Nabb et al., 2020). For Golgi-derived vesicles, kinesins likely bind at or near the TGN and we asked if KIF13A and KIF13B localized to vesicles in proximity of the TGN. We expressed Halo-KIF13A tail and stained for the cis-Golgi protein GM130 (Horton et al., 2005) and the trans-Golgi protein AP-1/γ1 (Farías et al., 2012; Jain et al., 2015) (Fig. 4A). High magnification images from a Golgi plane show that KIF13A colocalized with some AP-1/γ1 structures, but not with GM130. Colocalization quantification by Pearson’s correlation coefficient corroborated this observation (Fig. 4D). The correlation coefficient for GM130 and KIF13A was low (0.08) and substantially higher (0.28) for AP-1/γ1 and KIF13A. We performed equivalent experiments with Halo-KIF13B tail (Fig. 4B). KIF13B consistently labeled numerous structures that were proximal but not identical to GM130-labeled cis-Golgi (correlation coefficient: 0.28) (Fig. 4E). In contrast, KIF13B colocalized extensively with AP-1/γ1-labeled TGN (correlation coefficient: 0.47). Finally, we tested Halo-KIF1A tail (Fig. 4C). KIF1A is a Kinesin-3 family member that mediates transport of Golgi-derived dense core granules in dendrites and axons (Lo et al., 2011) and dendritic transport of low-density lipoprotein receptor, but not dendrite-selective TfR transport (Jenkins et al., 2012). Unlike the KIF13s, KIF1A did not localize to the Golgi (correlation coefficients: GM130 = −0.05, AP-1/γ1 = 0.07) (Fig. 4F). This shows that TGN targeting is specific for KIF13A and KIF13B and not a feature of all kinesins that mediate Golgi-derived vesicle transport.

**Figure 4:**
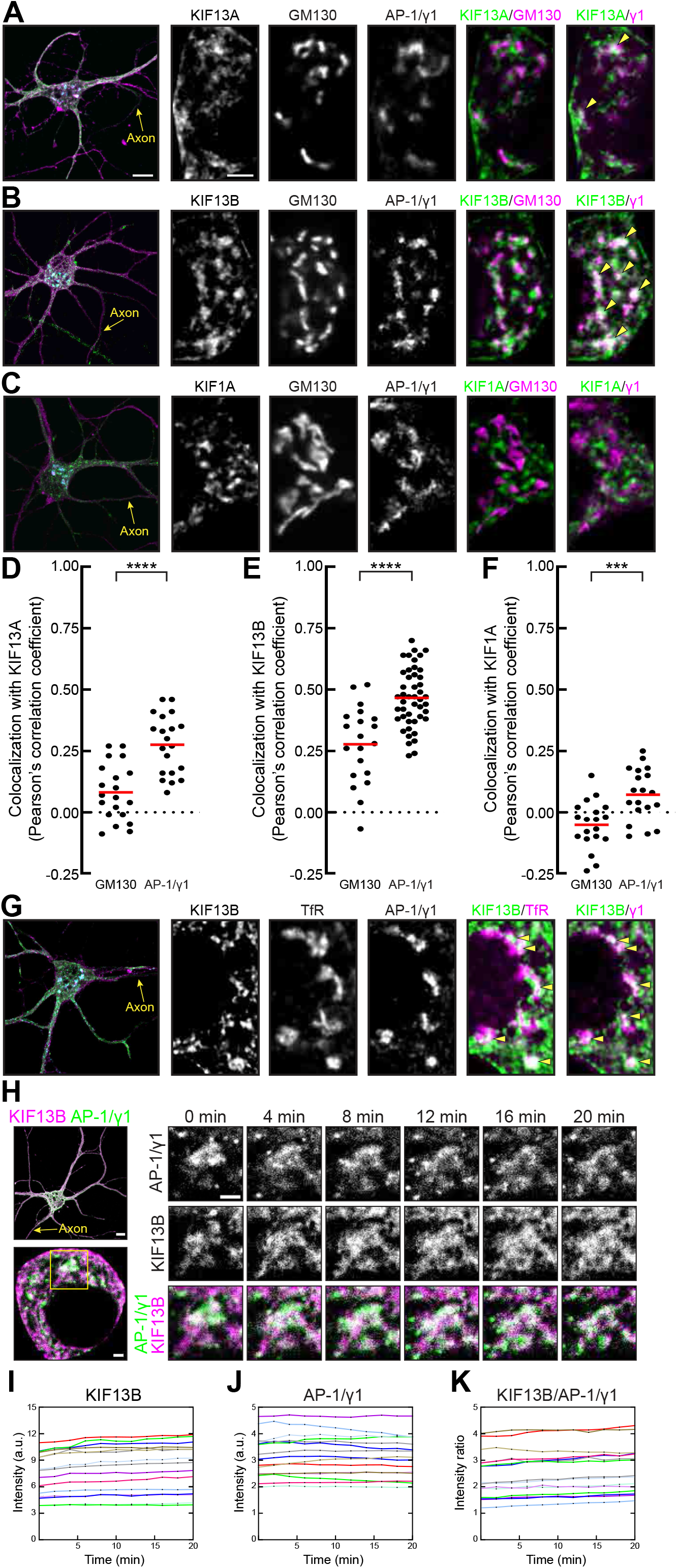
A stable KIF13B pool is maintained at the trans-Golgi network. (A-C&G) Representative images of 7-8 DIV hippocampal neurons expressing the indicated halo-tagged kinesin tail and stained for the indicated proteins. Halotags were visualized with JF549. High magnification images show a Golgi plane. Arrowheads indicate examples of colocalization. Scale bars: low magnification: 10 µm; high magnification: 3 µm. (D-F) Quantification of colocalization between indicated proteins in the Golgi plane. Cell numbers: KIF13A: 20; KIF13B/GM130: 20; KIF13B/γ1: 46; KIF1A: 19; ***p<0.001; ****p<0.0001. (H) Representative images of a 10 DIV hippocampal neuron expressing Halo-KIF13B tail and GFP-γ1. Halotag was visualized with JF549. Scale bars: low magnification: 10 µm; high magnification images: 2 µm. (I-K) Quantification of fluorescence intensity from a Golgi structure. Each line indicates measurements from one cell. Sample size: 15 cells.

Because KIF13B was enriched at the TGN and mediates some Golgi-derived TfR transport, we asked if TfR was present at KIF13B-positive TGN sites. We expressed Halo-KIF13B tail and immunolabeled endogenous TfR and AP-1/γ1 (Fig. 4G). TfR colocalized with AP-1/γ1 and KIF13B. Together, these data are the first evidence of kinesins being targeted to the mammalian TGN. They show that KIF13A and KIF13B are enriched at the TGN, possibly as a mechanism to ensure nascent vesicles carry the proper kinesin for transport immediately after budding.

Neurons are large cells that require a secretory pathway with the biosynthetic capacity to supply sufficient protein material (Futerman and Banker, 1996). That includes a dynamic TGN to sort cargoes into nascent transport vesicles (Ramazanov et al., 2021). We asked if KIF13B localization to the TGN was transient for individual budding events, or if KIF13B persisted at TGN sites dedicated to polarized vesicle budding. We coexpressed GFP-AP-1/γ1 to visualize TGN and Halo-KIF13B tail and generated 20 min live-cell recordings of a Golgi plane (Fig. 4H). In that time frame, most AP-1/γ1-labeled TGN sites persisted while undergoing some morphological dynamics. Each AP-1/γ1-labeled TGN site maintained a stable KIF13B pool during recording. Fluorescence Quantification of TGN sites from 15 neurons showed that the absolute amounts of KIF13B (Fig. 4I) and AP-1/γ1 (Fig. 4J) remained steady throughout recording. The relative distributions of KIF13B to AP-1/γ1 also remained stable, indicated by an unchanging intensity ratio (Fig. 4K). These results show that KIF13B-positive TGN structures are persistent, suggesting that they are sites specialized for the budding of polarized vesicles. This is the first example of any kinesin being stably targeted to the TGN of mammalian neurons and is evidence that KIF13B is maintained at the TGN instead of binding vesicles after budding.

### KIF13A and KIF13B bind AP-1/β1 in neurons

While the pool of KIF13s at the TGN was a surprising discovery, it provided the opportunity to determine the mechanistic link between vesicle budding and kinesin recruitment. We hypothesized that AP-1 may be part of that mechanistic link, because it mediates budding of dendrite-selective vesicles (Farías et al., 2012) and because classic yeast-two hybrid experiments found interaction between KIF13A and the ear domain of AP-1/β1 (Nakagawa et al., 2000). The AP-1/β1 ear is a small globular domain that is available in the cytoplasm when the AP-1 tetramer is bound to membranes (Bonifacino, 2014). It is distinct from the AP-1/β1 trunk domain which forms a tetrameric complex with other AP-1 subunits (Bonifacino, 2014; Pearse and Robinson, 1990), and the hinge domain that mediates binding to clathrin (Shih et al., 1995; Wilde and Brodsky, 1996). However, binding of KIF13A to AP-1 has not been tested in neurons and there is no evidence that KIF13B binds AP-1. While the motor domains of KIF13A and KIF13B are highly homologous, their vesicle-binding tails differ significantly. Therefore, binding of a protein to one KIF13 may not translate into equivalent binding of that protein to the other KIF13. To determine interactions between the ear domain of AP-1/β1 and KIF13s, we turned to a microscopy-based assay we developed for detecting protein–protein binding in their native milieu, the cytoplasm of intact neurons (Garbouchian et al., 2022). The assay uses two protein domains, the FK506 binding protein (FKBP) and FKBP12-rapmycin binding domain (FRB) that heterodimerize when cells are treated with a membrane permeant rapamycin analog (Banaszynski et al., 2005; Kapitein et al., 2010; Robinson et al., 2010; Jenkins et al., 2012; Bentley et al., 2015; Bentley and Banker, 2015; Belshaw et al., 1996). We coexpressed an FRB-tagged kinesin tail, the peroxisomal protein PEX3 fused to tdTomato and FKBP (PEX-tdTM-FKBP), and the GFP-tagged AP-1/β1 ear domain (GFP-β1^ear^) (Fig. 5A). Neurons were treated with or without linker for approximately 2 h before fixation and imaging. Linker-induced colocalization of GFP-β1^ear^ with peroxisomes indicates specific interaction between the candidate kinesin tail and the AP-1/β1 ear domain.

**Figure 5:**
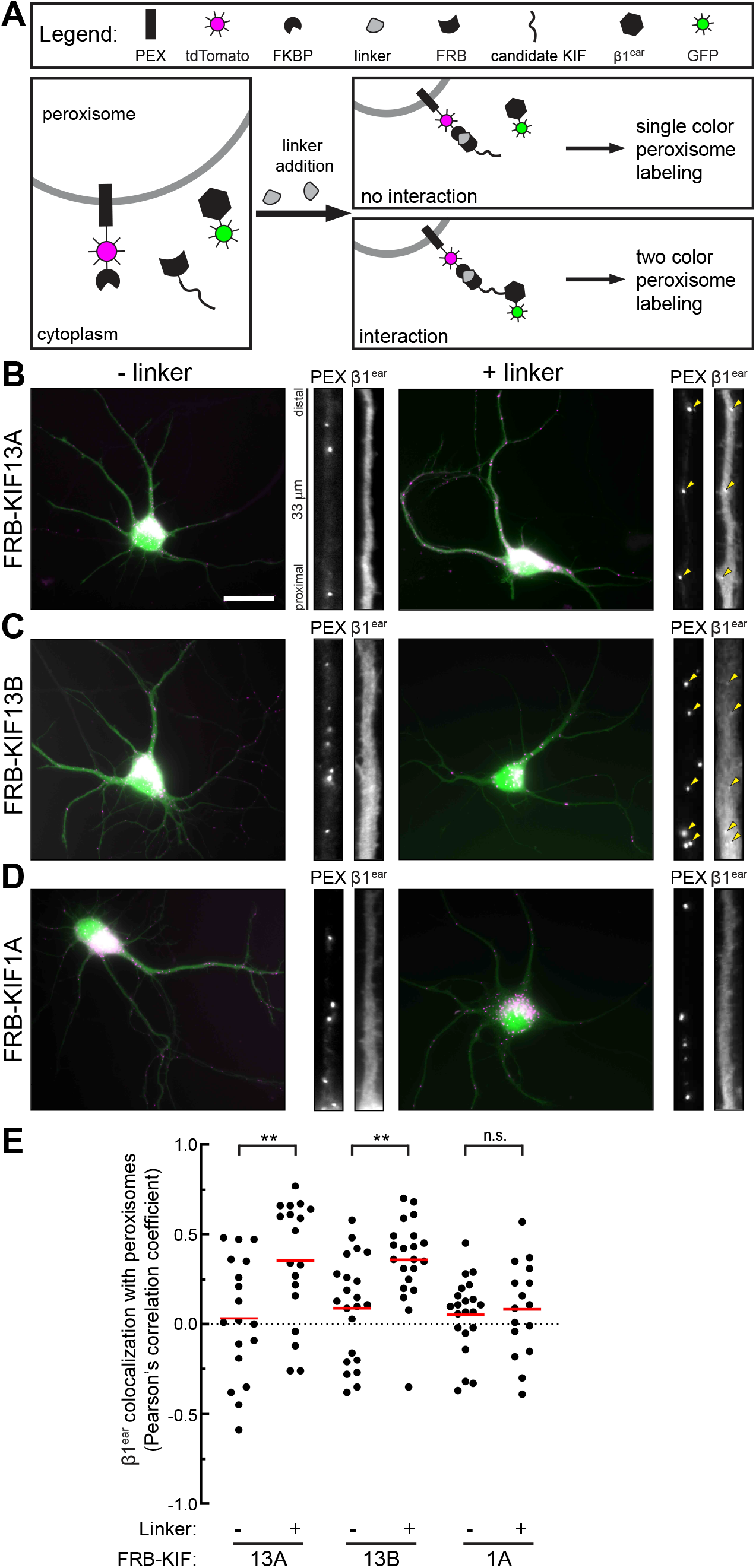
KIF13A and KIF13B bind AP-1/β1 in neurons. (A) Schematic of the protein–protein interaction assay. Three constructs are coexpressed: PEX-tdTomato-FKBP, GFP-β1^ear^, and an FRB-tagged candidate kinesin tail. Binding is evaluated by linker-induced colocalization of β1^ear^ and peroxisomes. (B-D) Representative images of 6-7 DIV hippocampal neurons expressing the indicated constructs. High magnification images from a representative dendrite show each channel separately. Arrowheads indicate examples of linker-induced colocalization. Scale bar: 20 µm. (E) Quantification of colocalization between peroxisomes and β1^ear^. Sample size: KIF13A(-): 18; KIF13A(+): 18; KIF13B(-): 22; KIF13B(+): 21; KIF1A(-): 21; KIF1A(+): 16. **p<0.01.

We first tested KIF13A (Fig. 5B). High magnification images from a dendrite show that PEX-tdTM-FKBP localized to peroxisomes as expected. GFP-β1^ear^ was soluble without any specific organelle labeling. In linker-treated neurons, GFP-β1^ear^ became specifically enriched on peroxisomes. We quantified colocalization by Person’s correlation coefficient (Fig. 5E). Without linker, the correlation coefficient was 0.03. In linker-treated neurons, the correlation coefficient increased to 0.35. This shows that KIF13A binds to the AP-1/β1 ear domain in neurons.

We used the same approach to test if KIF13B tail interacted with the AP-1/β1 ear domain (Fig. 5C). Without linker, GFP-β1^ear^ was soluble and produced a correlation coefficient of 0.09 (Fig. 5E). Linker treatment resulted in colocalization of GFP-β1^ear^ and peroxisomes that resulted in a correlation coefficient of 0.36. These data show conclusively that KIF13B tail binds the AP-1/β1 ear domain.

To determine if the interactions with the AP-1/β1 ear domain were specific for the kinesins that bind the TGN, we tested KIF1A in the same assay (Fig. 5D). Here linker treatment did not result in colocalization of GFP-β1^ear^ with peroxisomes. This was quantitatively shown by statistically indistinguishable correlation coefficients of plus- and minus-linker conditions (Fig. 5E).

These experiments show that both Kinesin-3 family members that are enriched at the TGN also interact with AP-1/β1, whereas KIF1A does not. The results argue that AP-1 maintains the pool of KIF13s at the TGN. This suggests a potential mechanism where polarized vesicles for which budding is mediated by AP-1 are automatically equipped with the proper kinesin. Therefore, AP-1 is likely to be the molecular link between the sorting of dendritically polarized cargoes into nascent vesicles and the recruitment of the proper kinesin motors for polarized transport.

### AP-1 colocalizes with axonal and dendritic cargoes at the TGN

While AP-1 is known to mediate sorting of dendritically polarized cargoes (Farías et al., 2012), the sorting machinery for axonally polarized cargo is unknown (Bentley and Banker, 2016; Guardia et al., 2018). In *C. elegans*, two different heterotetrameric clathrin adaptor complexes sort dendritically (AP-1) and axonally (AP-3) polarized membrane proteins (Li et al., 2016). AP-1 and AP-3 localize to distinct subdomains of the *C. elegans* TGN that are specialized for budding of dendritic and axonal vesicles (Li et al., 2016). To determine if mammalian neurons maintain comparable specialized TGN subdomains, we immunostained for AP-1/γ1 in neurons expressing either TfR-GFP or NgCAM-GFP (Fig. 6A&B) (Silverman et al., 2001; Wisco et al., 2003; Nabb and Bentley, 2022; Jareb and Banker, 1997; Hoo et al., 2016; Lewis et al., 2011). We quantified colocalization of each cargo with AP-1 in a Golgi plane and found that both colocalized equally with AP-1 (Fig. 6C). As a complementary approach, we performed immunostaining for endogenous AP-1/γ1, TfR, and L1CAM (the mammalian orthologue of NgCAM) in the same neurons and found AP-1/γ1 colocalization with both proteins (Fig. 6D). L1CAM is highly polarized to axons, and bright axonal staining is visible in the low magnification image. Measurement of correlation coefficients showed that AP-1/γ1 colocalized equally with both TfR (0.33) and L1CAM (0.35) (Fig. 6E). The absolute correlation coefficients in this experiment were lower than with exogenous proteins, likely due to lower signal-to-noise. However, the fact that colocalization of dendritic and axonal cargo with AP-1/γ1 was the same in both experiments, argues that previous observations with fluorescently tagged cargo proteins were not caused by overexpression effects. These experiments provide strong evidence that mammalian neurons do not have AP-1- specific TGN domains specialized for dendrite-selective vesicle budding, as exist in *C. elegans* (Li et al., 2016). Furthermore, these results suggest the possibility that AP-1 sorts both dendritic and axonal proteins in mammalian neurons.

**Figure 6:**
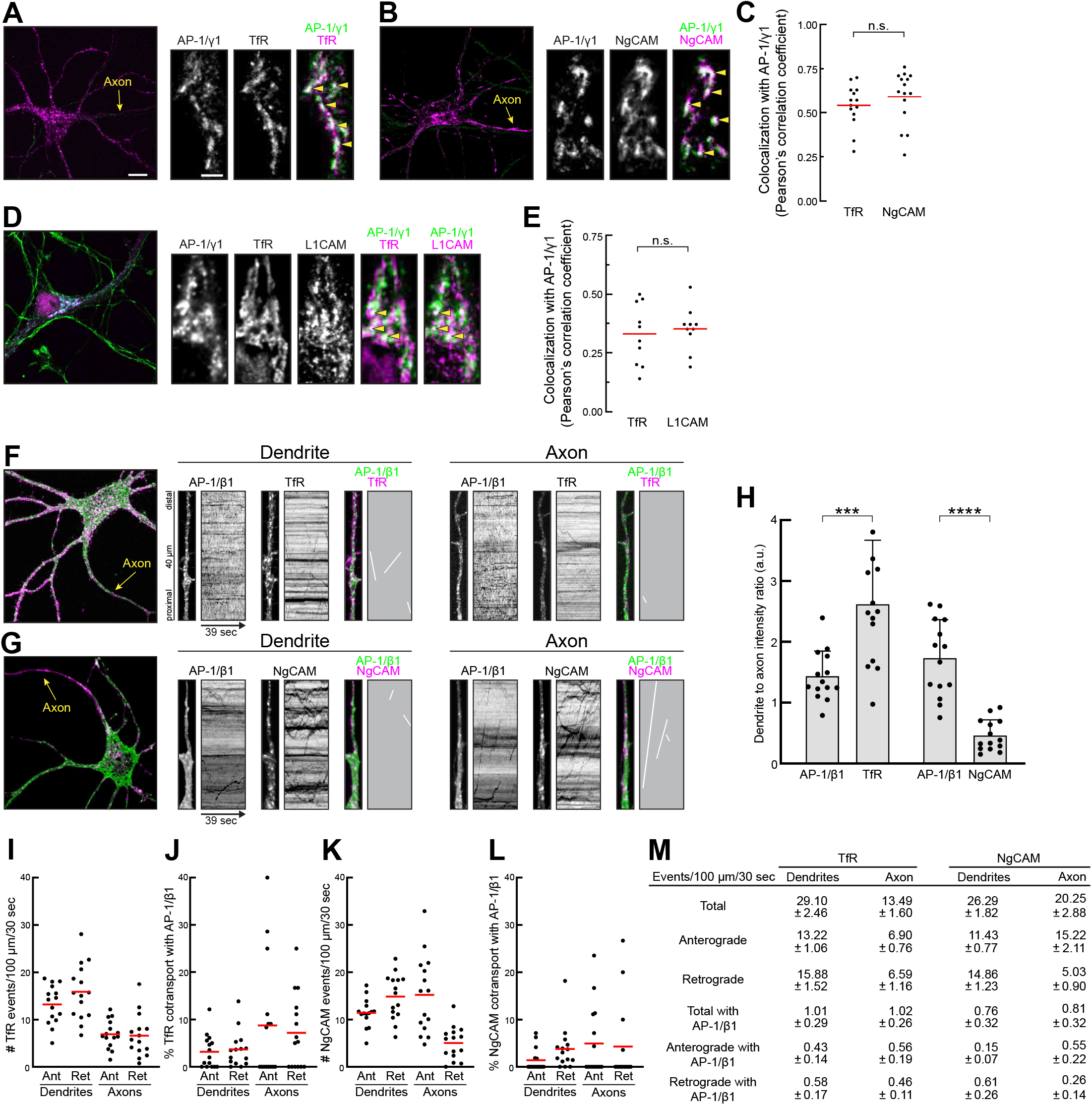
AP-1 colocalizes with dendritic and axonal cargoes at the TGN but is not a kinesin adaptor for long-range transport. (A&B) Representative images of 6 DIV hippocampal neurons expressing TfR-GFP (A) or NgCAM-GFP (B) and stained for AP-1/γ1. High magnification images show a representative Golgi plane. Arrowheads indicate examples of colocalization. Scale bars: low magnification: 10 µm; high magnification: 3 µm. (C) Quantification of colocalization. Cell numbers: TfR: 14; NgCAM: 15. (D) Representative image of a 9 DIV neuron stained for AP-1/ γ1, TfR, and L1CAM. High magnification images show a Golgi plane with arrowheads indicating examples of colocalization. (E) Quantification of AP-1/γ1 colocalization with TfR or L1CAM at the Golgi. Cell number: 10. (F&G) Representative images and kymographs of 7-8 DIV hippocampal neurons expressing AP-1/β1-Halo with TfR-GFP (F) or NgCAM-GFP (G). Halotags were visualized with JF549. (H) Quantification of dendrite to axon ratios for each protein. One TfR data point was omitted because it exceeded the y-axis. Cell counts: TfR: 14; NgCAM: 14. ***p<0.001; ****p<0.0001. (I-L) Quantification of TfR and NgCAM transport events and cotransport of each with AP-1/β1 in the axon and dendrites. (M) Table displaying event and cotransport frequency for each condition. (I-M) Sample size: TfR: 15 cells; NgCAM: 15 cells.

### AP-1 is not the long-term kinesin adaptor for polarized vesicle transport

The function of heterotetrameric clathrin adaptor complexes during cargo sorting and vesicle budding are well established (Bonifacino, 2014; Duncan, 2022; Dell’Angelica and Bonifacino, 2019). It is less clear what happens with AP complexes after budding. AP complexes are thought to dissociate relatively soon after vesicle budding completes (Bonifacino, 2014; Sanger et al., 2019; Dell’Angelica and Bonifacino, 2019), although some studies suggest that AP complexes can act as long-term motor adaptors on vesicles (Horikawa et al., 2002; Schmidt et al., 2009; Azevedo et al., 2009; Maritzen and Haucke, 2010), including for KIF13A in non-neuronal cells (Nakagawa et al., 2000; Delevoye et al., 2009; Campagne et al., 2018).

To determine if AP-1 is the transport adaptor for KIF13s, we asked if the cargoes that colocalized with AP-1 at the TGN also cotransported with AP-1. We coexpressed Halo-tagged AP-1/β1 with either TfR-GFP or NgCAM-GFP and performed live-cell imaging (Fig. 6F&G). TfR and NgCAM exhibited the anticipated dendritic and axonal polarities (Fig. 6H). Kymographs show that TfR and NgCAM exhibited the expected dendrite- or axon-selective transport behaviors (Fig. 6F&G). High magnification images identify AP-1-labeled vesicles in dendrites and axons and the associated kymographs show that most AP-1 vesicles were stationary. Few of the stationary vesicles colocalized with TfR or NgCAM. We then quantified cotransport of TfR or NgCAM with AP-1/β1 (Fig. 6J&L). In all conditions cotransport was minimal. The only condition exceeding 8 % cotransport was for axonal TfR, where TfR events were rare. It is likely that these are wayward TfR vesicles recently retrieved from the plasma membrane by endocytosis. Fig. 6M shows total event numbers for TfR and NgCAM vesicles and minimal cotransport with AP-1 in each condition. A potential caveat of this experiment is that labeling vesicles with AP-1 is challenging and may miss some cotransport. However, the fact that we observed substantial AP-1 labeling of stationary organelles suggests that this was not the reason for little cotransport. Therefore, the small amount of vesicle cotransport with AP-1 and the fact that most AP-1-labeled vesicles were stationary, are strong evidence that AP-1 is not the long-term kinesin adaptor for these vesicles. Instead, these results point to AP-1 maintaining the TGN pool of KIF13s. There may be a “handoff” to other proteins that become the kinesin adaptors for long-range vesicle transport after or during budding.

### Interaction between AP-1 and KIF13s is required for proper polarized vesicle transport

Taken together, the data suggest the hypothesis that AP-1 plays a central role in recruiting KIF13s to both dendrite- and axon-selective vesicles. To test this hypothesis, we designed a dominant-negative approach that specifically interfered with KIF13 binding to AP-1, without interfering with other functions of endogenous AP-1. Because β1^ear^ mediates KIF13B binding (Fig. 5) its overexpression may saturate AP-1 binding sites on KIF13A and KIF13B without impacting the cargo sorting and TGN budding activity of endogenous AP-1/β1.

We treated neurons expressing GFP fill or GFP-β1^ear^ for 48 h with Tf for 30 min (Fig. 7A&B). If KIF13s transport newly synthesized TfR to the dendritic membrane and this function depends on binding to AP-1, expression of GFP-β1^ear^ would decrease TfR at the membrane and reduce Tf uptake. The total amount of Tf uptake was reduced in neurons expressing GFP-β1^ear^. Measurement of Tf fluorescence in dendrites found a significant decrease in GFP-β1^ear^- expressing neurons (Fig. 7C). Measuring the number of Tf-labeled endosomes found no reduction in endosomes (Fig. 7D). Therefore, expression of GFP-β1^ear^ does not impact the frequency of clathrin-mediated endocytosis. This interpretation is strengthened by the fact that Tf did not build up in the dendritic plasma membrane, as it would if GFP-β1^ear^ interfered with clathrin-mediated endocytosis of endogenous transferrin receptors. Instead, decreased Tf uptake without decreased endocytosis shows that fewer transferrin receptors were available to import fluorescent transferrin for each endocytic event. This supports the model that KIF13–AP-1 interactions are required for proper transport of TfR to the plasma membrane.

**Figure 7:**
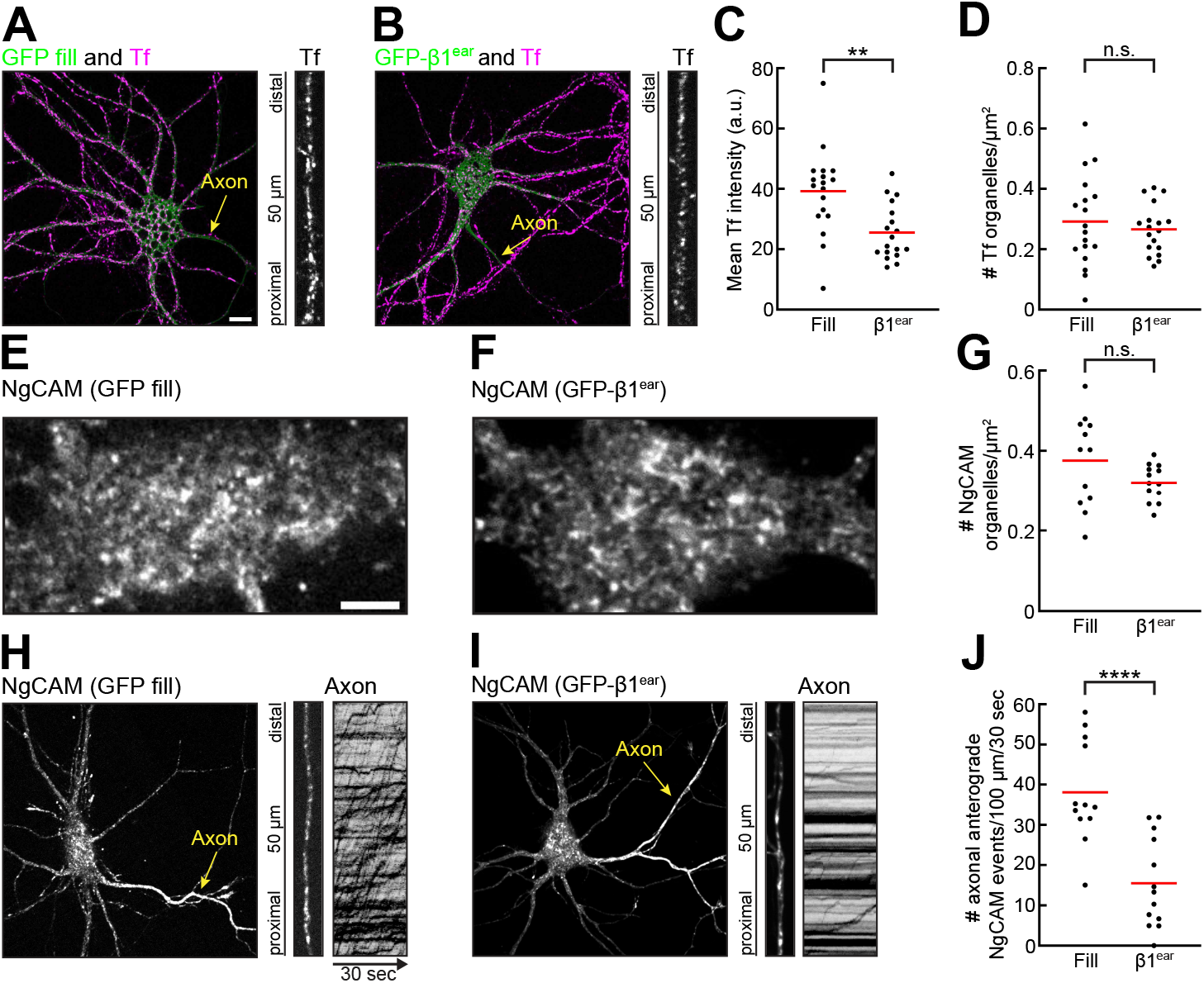
AP-1 and KIF13B interactions are required for proper Tf uptake and axon-selective transport. (A&B) Representative images of 9 DIV hippocampal neurons expressing (A) GFP fill or (B) GFP β1^ear^ for 48 h and incubated with Tf647 for 30 min. Scale bar: 10 µm. (C) Quantification of Tf fluorescence intensity in dendrites. (D) Quantification of Tf vesicles in the same region as (C). Cell counts: GFP fill: 18; GFP-β1^ear^: 18. **p<0.01. (E&F) high magnification images of the somata of 10 DIV hippocampal neurons expressing NgCAM-Halo with either (E) GFP fill or (F) GFP-β1^ear^. Scale bar: 5 µm. (G) Quantification of NgCAM vesicles in somata. (H&I) Representative images and kymographs of neurons from (E&F). Transfected neurons were incubated with JF549 for 15 m to saturate existing NgCAM-Halo binding sites. Two hours after JF549 washout, cells were treated with JF646 and imaged. (J) Quantification depicting anterograde NgCAM events in axons. Sample size: GFP fill: 12 cells; GFP-β1^ear^: 13 cells. ****p<0.0001.

Because KIF13B mediates the transport of axon-selective NgCAM vesicles (Fig. 3), we asked if KIF13B–AP-1 interactions were required for axonal transport of NgCAM. Long-term expression of NgCAM results in its buildup in the axonal membrane (Sampo et al., 2003; Nabb and Bentley, 2022), which prevents imaging of vesicles in the axon. To visualize axonal NgCAM transport, we used the halotag system (Los et al., 2008; Encell, 2012). The halotag enzyme itself is not fluorescent. Fluorescence is conferred by covalent binding to a ligand with an organic dye. Because enzyme–substrate binding is covalent, binding of a ligand is irreversible which prevents the halotag enzyme from binding further ligands. Sequential exposure to spectrally distinct ligands can specifically visualize newly synthesized proteins (Yoon et al., 2016; Frank et al., 2022).

We coexpressed NgCAM-Halo with a GFP fill or GFP-β1^ear^ for 48 h. Two hours prior to imaging, neurons were treated with JF549 for 15 min. Before imaging, neurons were incubated with JF646 to visualize NgCAM that has been synthesized in the 2 h since JF549 exposure. This treatment resulted in NgCAM vesicle labeling equivalent to that observed after typical short-term NgCAM expression (Nabb and Bentley, 2022; Frank et al., 2022). We first asked if GFP-β1^ear^ expression reduced vesicle budding at the TGN. If budding was affected there would be fewer vesicles in the somata. We measured the number of vesicles containing newly synthesized NgCAM in non-Golgi planes of somata (Fig. 7E&F). There was no difference in the vesicle numbers between neurons expressing GFP fill and those expressing GFP-β1^ear^ (Fig. 7G). This was further supported by the fact that newly synthesized NgCAM still reached the axon in the two hours after JF549 treatment (Fig. 7H&I). Therefore, budding of NgCAM vesicles from the TGN was not affected by GFP-β1^ear^ expression.

We examined NgCAM vesicle transport in the axon by kymograph analysis. Neurons expressing GFP-β1^ear^ exhibited substantially less NgCAM transport in axons than those expressing a GFP fill (Fig. 7H&I). Quantification of anterograde transport events found that axonal NgCAM transport in neurons expressing GFP-β1^ear^ was decreased significantly, dropping from 38 events/100 µm/30 sec in GFP-expressing cells to 16 events/100 µm/30 sec in cells with GFP- β1^ear^ (Fig. 7J). This decrease in transport shows that interactions between KIF13B and AP-1/β1 are required for the transport of axon-selective vesicles.

Together, these data show that interactions between AP-1 and KIF13s are crucial for trafficking of both dendritically and axonally polarized membrane proteins. Because AP-1 is crucial for TGN sorting, these data also support the model that AP-1 is the molecular link between polarized cargo sorting and subsequent kinesin recruitment.

## DISCUSSION

In this study, we defined the functions of Kinesin-3 family members KIF13A and KIF13B in polarized vesicle transport. We found that KIF13A is a dedicated dendrite-selective kinesin and that KIF13B mediates transport of both dendrite- and axon-selective vesicles. Neurons maintain a stable pool of KIF13B and, to a lesser extent, KIF13A at the TGN. Polarized vesicles originate at the TGN (Farías et al., 2012; Nabb and Bentley, 2022) and this pool provides a convenient kinesin source for nascent vesicles. KIF13A and KIF13B bind AP-1, the adaptor complex that mediates the sorting of polarized vesicles at the TGN. Interference with AP-1–KIF13 interactions disrupted delivery of dendrite-selective membrane proteins and reduced axonal transport of axon-selective vesicles. Together, these data suggest a new model for KIF13 activity in polarized trafficking (Fig. 8). In this model, KIF13A and KIF13B are maintained at the TGN by interactions with AP-1. Interaction with AP-1 recruits KIF13s specifically to Golgi-derived vesicles destined for polarized transport. KIF13A exclusively participates in the transport of dendrite-selective vesicles; those that move from the TGN to the plasma membrane as well as that of dendrite-selective endosomes. KIF13B performs dendrite-selective transport and plays an additional role in axon-selective transport. This model defines the specific transport steps that are mediated by KIF13A and KIF13B and defines the trafficking framework for future studies to determine the molecular mechanisms by which KIF13A and KIF13B confer selective transport.

**Figure 8:**
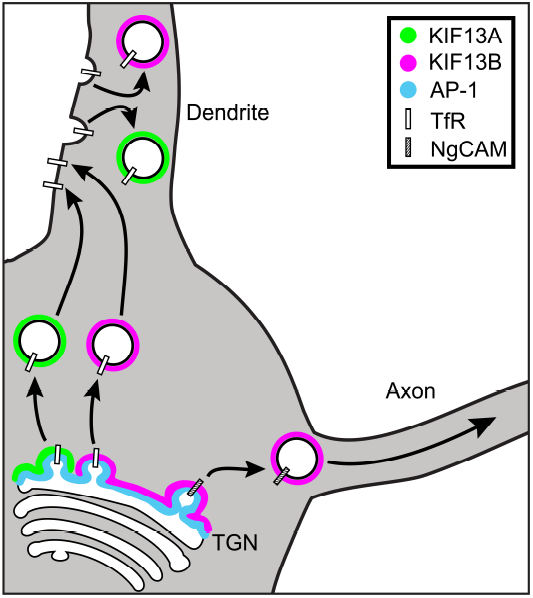
KIF13A and KIF13B perform distinct functions in polarized transport. KIF13A is targeted to a subset of the TGN and mediates dendrite-selective transport of Golgi-derived and endocytic vesicles. KIF13B and, to a lesser degree, KIF13A are enriched at the TGN by binding AP-1. KIF13A performs dendrite-selective transport of Golgi-derived and endocytic vesicles. KIF13B can perform dendrite-selective transport, but also mediates a second function by transporting axon-selective vesicles.

### Mechanisms of dendrite-selective transport and on-vesicle regulation of KIF13s

Vesicles that contain dendritically polarized membrane proteins undergo long-range movement in dendrites, but suspend their movement in the proximal axon (Petersen et al., 2014). The mechanisms by which vesicles are arrested in the proximal axon have gained significant attention. Despite this, no comprehensive mechanistic model has emerged (Bentley and Banker, 2016; Radler et al., 2020). Most studies proposed models in which dendrite-selective vesicles are “captured” by mechanisms that are external to the vesicles. Examples include myosin V-mediated capture on actin filaments in the proximal axon (Al-Bassam et al., 2012; Watanabe et al., 2012; Janssen et al., 2017; Lewis et al., 2009), dynein-mediated retrograde movement that clears dendritic vesicles from the axon (Kapitein et al., 2010; Kuijpers et al., 2016; Janssen et al., 2016), and membrane fusion of polarized vesicles in the proximal axon (Eichel et al., 2022). These models are “kinesin agnostic”, in that the identity of the kinesins that move dendrite-selective vesicles are not part of the mechanism that confers dendritic selectivity. Little attention has been paid to the kinesin motors that transport dendrite-selective vesicles. However, kinesin regulation may contribute to dendritic selectivity as general evidence for on-vesicle regulation of kinesins has grown steadily (Wang and Schwarz, 2009; Fu and Holzbaur, 2013; Kelliher et al., 2018; Yang et al., 2019; Keren-Kaplan and Bonifacino, 2021). The fact that two closely related kinesins— KIF13A and KIF13B—seem to mediate most dendrite-selective transport may hint at the existence of a common, kinesin-focused mechanism for dendrite-selective transport.

On-vesicle regulation is a growing theme in kinesin-mediated vesicle transport (Nabb et al., 2020). KIF13B may be a useful model kinesin for on-vesicle regulation, because our data corroborate and extend previous studies that proposed KIF13B mediates dendrite-selective transport (Jenkins et al., 2012) while also participating in axon selection and extension (Horiguchi et al., 2006; Yoshimura et al., 2010). A likely source of transport regulation are the vesicle adaptors that target kinesins to specific vesicles. Pioneering studies from Chishti and colleagues identified hDdlg/SAP97 as a KIF13B regulatory protein (Hanada et al., 2000; Asaba et al., 2003; Yamada et al., 2007; Mills et al., 2019). SAP97 is thought to recruit KIF13B to vesicles and activate motor activity. Because SAP97 mediates trafficking of dendritic AMPA receptors (Sans et al., 2001; Rumbaugh et al., 2003; Waites et al., 2009; Zheng and Keifer, 2014; Goodman et al., 2017), it is a promising candidate to participate in KIF13B-mediated dendrite-selective transport. A candidate regulatory protein for axon-selective KIF13B transport is the Arf6 GAP centaurin-α1. Centaurin-α1 binds the FHA domain of KIF13B (Venkateswarlu et al., 2005; Tong et al., 2010), which may be part of the mechanism by which KIF13B is recruited to axon-selective vesicles. Future studies will need to integrate these fragments within the overall framework of neuronal KIF13B trafficking.

### The molecular link between cargo sorting and kinesin recruitment

Cargo sorting and the mechanics of vesicle transport have typically been studied as independent phenomena but must be linked in cells to ensure that cargo proteins reach the correct destination. This study provides the first evidence that AP-1 connects sorting of polarized cargoes into vesicles with the recruitment of specific kinesins. For dendritically polarized vesicles, evidence is converging on a clear model. AP-1 maintains KIF13A and KIF13B at the TGN. During vesicle formation, AP-1 recognizes dendritically polarized cargoes at the TGN by a tyrosine- or dileucine-based motif and packages them into vesicles destined for dendrite-selective transport (Farías et al., 2012; Jain et al., 2015). Despite this progress, significant questions remain. AP-1 is thought to dissociate from vesicles soon after budding (Ghosh and Kornfeld, 2003), which is consistent with the observation that few polarized vesicles cotransported with AP-1. These data argue strongly that AP-1 is not the long-term kinesin adaptor for polarized vesicles, despite mediating the initial recruitment of kinesins to the TGN. The mechanism by which KIF13s are “handed off” from AP-1 to the long-term kinesin adaptor is unclear; at present, the identity of the long-term kinesin adaptor is unknown. Furthermore, the mechanism by which dendritically polarized proteins are recognized is not fully understood. The AP-1/µ1 subunits binds some dendritically polarized proteins by a YXXΦ motif. However, the YXXΦ motif alone cannot be sufficient for specific dendritic sorting, as many membrane proteins possess compatible sequences, but most do not exhibit dendritic polarity. For example, axonally polarized NgCAM has a canonical YXXΦ that is required for AP-2-mediated endocytosis in axons (Schaefer et al., 2002; Kamiguchi et al., 1998). This suggests the existence of secondary signals, which remain to be identified.

The sorting and transport of axonally polarized proteins is even less understood. Classic experiments evaluating axon-selective sorting failed to identify any consensus sorting motifs (Sampo et al., 2003; Yap et al., 2008) and there has been little progress since (Bonifacino, 2014; Bentley and Banker, 2016). We found that disruptions to interactions between KIF13B and AP-1 decreased axonal transport. This result suggests that AP-1 participates in axonal sorting. If AP-1 mediates the TGN sorting of both dendritically and axonally polarized proteins, some yet unknown factor mediates the separation of these proteins into different Golgi-derived vesicles. This mechanism appears to be efficient, as there is little cotransport of dendritically and axonally polarized membrane proteins (Petersen et al., 2014).

KIF13B plays a previously unappreciated role in axon-selective transport. Current dogma argues that Kinesin-1 family members (KIF5A, KIF5B, and KIF5C) are the primary axon-selective motors (Nabb et al., 2020). Kinesin-1 was first identified from axoplasm (Vale et al., 1985b; a) and constitutively active Kinesin-1 motor domains accumulate in axon tips (Nakata and Hirokawa, 2003; Jacobson et al., 2006; Huang and Banker, 2012). Furthermore, Kinesin-1 mediates transport of multiple axonal vesicles, including lysosomes (Farías et al., 2017), mitochondria (Wang and Schwarz, 2009), amyloid precursor protein vesicles (Fu and Holzbaur, 2013), and GPI-anchored protein (Encalada et al., 2011). Together, these observations led to the assumption that Kinesin-1 family members also mediate axon-selective transport (Bentley and Banker, 2016; Nabb et al., 2020). The substantial axonal cotransport of KIF13B and NgCAM suggest a prominent role for KIF13B in axon-selective transport. This does not necessarily mean that KIF13B is the exclusive axon-selective kinesin. One possibility is that Kinesin-1 and KIF13B cooperate in axon-selective transport. Alternatively, Kinesin-1 and KIF13B may move different axon-selective subpopulations, with NgCAM vesicles being only one of several.

## MATERIALS AND METHODS

### Cell culture

Primary hippocampal neurons were cultured as previously described (Kaech and Banker, 2006; Kaech et al., 2012b). E18 rat hippocampi were dissected, trypsinized, dissociated, and plated onto 18-mm glass coverslips coated with poly-l-lysine. Cultured neurons were grown in N2- supplemented MEM and maintained at 37°C with 5% CO2. Stage 4 hippocampal neurons (5–10 DIV) were transfected with Lipofectamine 2000 (Thermo Fisher, Cat# 11668019). The following expression times were used: Figs. 1-3, 6A-C: 16-20 h; Fig. 4A-G: 5-8 h; Fig. 5: 12 h; Figs. 4H-K, 6F-M: 24 h; Fig. 7: 48 h.

For most immunofluorescence experiments, neurons were fixed with 4 % paraformaldehyde (PFA)/4 % sucrose in phosphate-buffered saline (PBS) for 20-45 min at 37°C. PFA-treated neurons were permeabilized with 0.5 % Triton X-100 in PBS. For L1CAM staining (Fig. 6D&E), neurons were fixed with methanol supplemented with 5 mM EGTA for 11-20 min at −20°C. Fixed neurons were blocked with 0.5 % fish skin gelatin in PBS for at least 1 h. Coverslips were incubated with primary antibodies for at least 1 h, washed with PBS, incubated with secondary antibodies for at least 30 min, and mounted on slides with Prolong Diamond Anti-fade (Invitrogen). The following antibodies and dilutions were used in this study: AP-1/γ1-adaptin (1:1000, BD Biosciences, Cat#: 610385), GM130 (1:1000, BD Biosciences, Cat#: 610823), TfR (1:1000, ThermoFisher, Cat#: 13-6800), L1CAM (1:1000, Abcam, Cat#: AB24345).

### DNA constructs

Construct details are described in Table 1. All constructs were confirmed by sequencing.

**Table 1:**
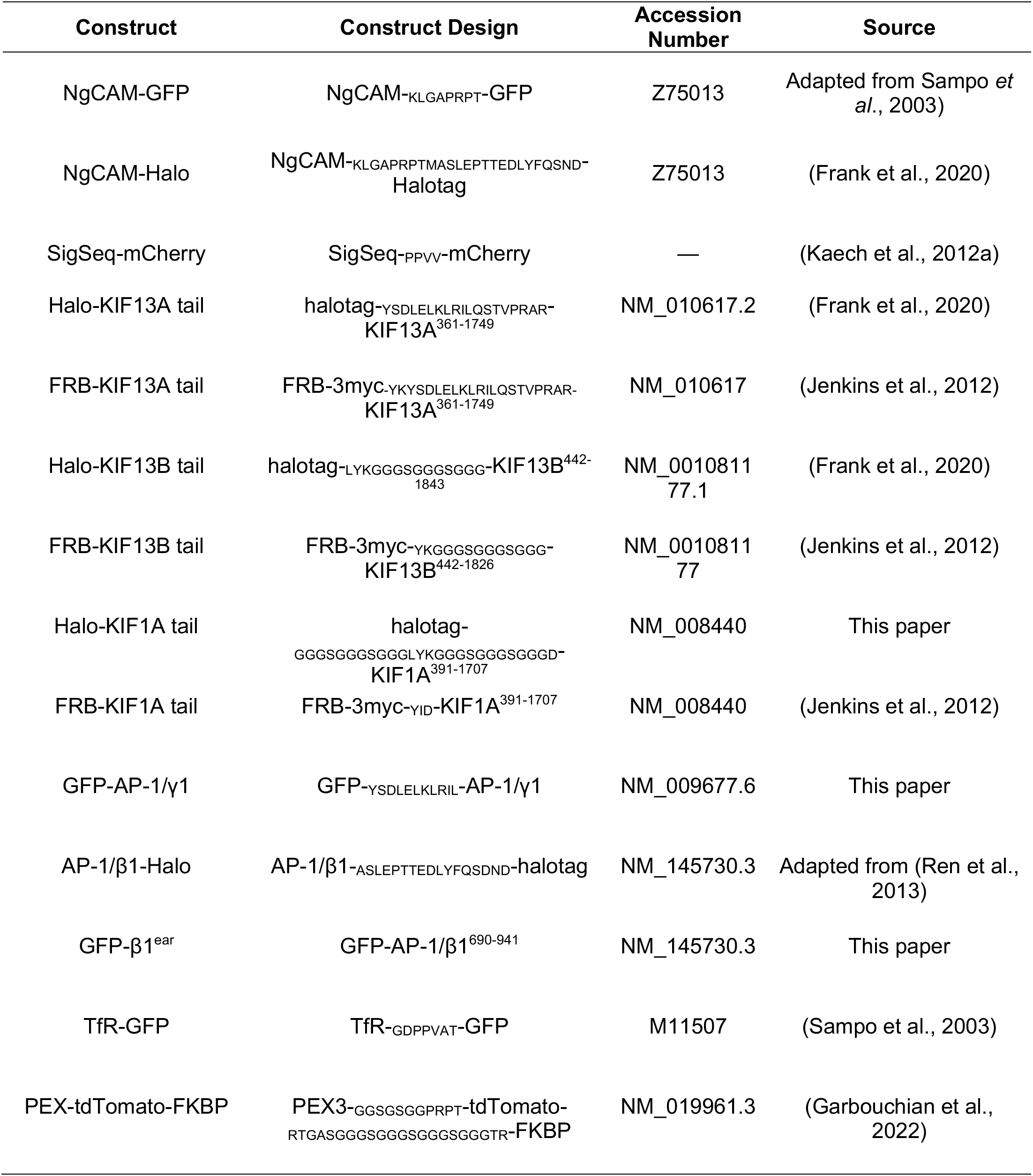
Expression constructs.

### Halo labeling, transferrin treatment, and peroxisome targeting

Halotags were visualized with JF549 or JF646 (Grimm et al., 2015) as indicated in the figure legends. Neurons were incubated with 10 nM JF dye for at least 5-15 minutes before imaging or fixation. For Fig. 7 experiments, NgCAM-Halo was expressed for 48 h and incubated with JF549 for 15 min. After JF washout and an additional 2 h incubation, cells were rinsed and incubated with JF646 for 5 min prior to imaging.

For transferrin labeling, neurons were incubated with 40 nM Tf555 or Tf647 (Thermo Fisher, Cat# T35352 and T23366) for at least 20 min prior to live cell imaging (Fig. 2) or 30 min prior to fixation (Fig. 7).

For Fig. 5, neurons expressing the indicated constructs were incubated with or without 100 nM linker (AP21967; Takara Cat#635056) for 2 h prior to fixation and imaging.

### Imaging

Most images and all movies were acquired with an Andor Dragonfly built on a Ti2 (Nikon) with a CFI Apo 60x 1.49 objective (Nikon) and two sCMOS cameras (Zyla 4.2+; Andor). The imaging stage, microscope objectives, and cell samples (live or fixed) were maintained at 37°C in a warmed enclosure (full lexan incubation ensemble; OkoLab). The z-axis movement was controlled by a Perfect Focus System (Nikon). Most movies were recorded at approximately 1.4 or 2 frames per second. Movies in Fig. 4H-K were recorded for 20 min at 0.2 frames per second. Axons were identified by morphology and polarity of constructs. Images in Fig. 5 were acquired with a Zeiss Axio Imager Z1 microscope (Carl Zeiss) equipped with an Axiocam 506 mono.

### Analysis

MetaMorph Image Analysis Software (Molecular Devices) was used to generate kymographs of vesicle transport. All kymograph analyses except Figs. 6F-M and 3 were performed by a single blinded analyst. For Fig. 1, only three dendrites were analyzed per cell. Transport events were identified on kymographs where each continuous line with a consistent slope was scored as a single transport event. A single vesicle could undergo multiple transport events if there was a distinct pause between them. Once all event coordinates had been identified, cotransporting events were determined by comparing the locations, slopes, and run lengths from both channels. Only events that demonstrated substantial overlap were scored. Event coordinates were exported to Microsoft Excel for analysis. Event velocities and run lengths (distance moved along the long axis of the neurite) were calculated. To ensure only microtubule-based long-range transport events were included in the analysis, excursions <3 μm were excluded. The cotransport percentage was calculated as the number of overlapping events divided by the total number of events. The proximal 25 µm of the axon were excluded for axonal analysis in Fig. 6F-M, due to the proximal axon possessing dendritic properties.

For Figs. 1&2 colocalization was determined by blinded manual object counting in ImageJ in the somata or dendrites, as indicated. Dendritic object counting was performed in one 20 µm section of one dendrite per cell.

Pearson’s correlation coefficients were calculated in ImageJ with the coloc2 plugin. Golgi colocalization (Figs. 4A-F, 6A-E) was calculated for a polygon in the soma that excluded the nucleus. Dendrite colocalization (Fig. 5) was calculated for a polygon that was drawn around 3-5 peroxisomes.

Dendrite to axon ratios were calculated by a single blinded analyst. In the first frame of each movie, a 20 pixel wide line scan was generated from the primary dendrite and the axon. The bottom 10th percentile of the mean intensity was subtracted to remove background. The dendrite to axon ratio was calculated by dividing average intensity of the first 20 μm of the primary dendrite by the average intensity of the axonal region 60-80 μm away from the soma (beyond the axon initial segment).

Tf object counting and intensity analyses (Fig. 7A-D) were performed blinded in 15–25 µm regions of an isolated dendrite from each neuron. For intensity measurements, a linear subtraction was performed to remove camera background. The same dendrite regions were used for both analyses. NgCAM object counting analysis (Fig. 7E-G) was performed blinded in the somata of transfected neurons.

Analysis of long-term TGN imaging (Fig. 4H-K) was performed in a circular ROI with a 40- pixel diameter that was drawn around a stable AP-1/γ1 structure. The absolute intensity of this region was measured in both channels at the indicated time points.

One-way ANOVA with Tukey multiple comparisons was used to evaluate differences in the dendrite to axon ratios between KIF13A, TfR, and SigSeq as well as between KIF13B, TfR, and SigSeq experiments. Student’s t-tests were used to determine P values for all the other compared conditions (visualized with brackets) using GraphPad Prism. All plots were generated in GraphPad Prism.

## ACKNOWLEDGEMENTS

We thank Drs. Catherine Drerup, Susan P. Gilbert, and Gary Banker and members of the Bentley lab for their helpful comments on the paper. We acknowledge the BioResearch facility at Rensselaer for assistance with husbandry and tissue collection.

## AUTHOR CONTRIBUTIONS

Conceptualization, project administration, and supervision: M. Bentley. Investigation: A.C. Montgomery, C.S. Mendoza, and G.B. Quinones. Data curation and analysis: A.C. Montgomery, C.S. Mendoza, and M. Bentley. Visualization: A.C. Montgomery, C.S. Mendoza, A. Garbouchian, and M. Bentley. Writing, original draft and editing: A.C. Montgomery and M. Bentley.

## Abbreviations

TGN: trans-Golgi network
AP: heterotetrameric clathrin adaptor protein complex
TfR: transferrin receptor
Tf: transferrin
NgCAM: neuron-glia cell adhesion molecule
FKBP: FK506 binding protein
FRB: FKBP12-rapamycin binding domain.

## REFERENCES

Al-Bassam, S., M. Xu, T.J. Wandless, and D.B. Arnold. 2012. Differential trafficking of transport vesicles contributes to the localization of dendritic proteins. Cell Rep. 2:89–100. doi:10.1016/j.celrep.2012.05.018.

Asaba, N., T. Hanada, A. Takeuchi, and A.H. Chishti. 2003. Direct interaction with a kinesin-related motor mediates transport of mammalian discs large tumor suppressor homologue in epithelial cells. J. Biol. Chem. 278:8395–8400. doi:10.1074/jbc.M210362200.

Azevedo, C., A. Burton, E. Ruiz-Mateos, M. Marsh, and A. Saiardi. 2009. Inositol pyrophosphate mediated pyrophosphorylation of AP3B1 regulates HIV-1 Gag release. Proc. Natl. Acad. Sci. U. S. A. 106:21161–21166. doi:10.1073/pnas.0909176106.

Banaszynski, L.A., C.W. Liu, and T.J. Wandless. 2005. Characterization of the FKBP-rapamycin-FRB ternary complex. J. Am. Chem. Soc. 127:4715–4721. doi:10.1021/ja043277y.

Belshaw, P.J., S.N. Ho, G.R. Crabtree, and S.L. Schreiber. 1996. Controlling protein association and subcellular localization with a synthetic ligand that induces heterodimerization of proteins. Proc. Natl. Acad. Sci. U. S. A. 93:4604–4607. doi:10.1073/pnas.93.10.4604.

Bentley, M., and G. Banker. 2015. A Novel Assay to Identify the Trafficking Proteins that Bind to Specific Vesicle Populations. Curr. Protoc. cell Biol. 69:13.8.1–13.8.12. doi:10.1002/0471143030.cb1308s69.

Bentley, M., and G. Banker. 2016. The cellular mechanisms that maintain neuronal polarity. Nat. Rev. Neurosci. 17:611–622. doi:10.1038/nrn.2016.100.

Bentley, M., H. Decker, J. Luisi, and G. Banker. 2015. A novel assay reveals preferential binding between Rabs, kinesins, and specific endosomal subpopulations. J. Cell Biol. 93:4604. doi:10.1083/jcb.201408056.

Bonifacino, J.S. 2014. Adaptor proteins involved in polarized sorting. J. Cell Biol. 204:7–17. doi:10.1083/jcb.201310021.

Burack, M.A., M.A. Silverman, and G. Banker. 2000. The role of selective transport in neuronal protein sorting. Neuron. 26:465–472. doi:10.1016/S0896-6273(00)81178-2.

Campagne, C., L. Ripoll, F. Gilles-Marsens, G. Raposo, and C. Delevoye. 2018. AP-1/KIF13A Blocking Peptides Impair Melanosome Maturation and Melanin Synthesis. Int. J. Mol. Sci. 19:568. doi:10.3390/ijms19020568.

Craig, A.M., and G. Banker. 1994. Neuronal polarity. Annu Rev Neurosci. 17:267–310. doi:10.1146/annurev.ne.17.030194.001411.

Das, U., D.A. Scott, A. Ganguly, E.H. Koo, Y. Tang, and S. Roy. 2013. Activity-induced convergence of app and bace-1 in acidic microdomains via an endocytosis-dependent pathway. Neuron. 79:447–460. doi:10.1016/j.neuron.2013.05.035.

Delevoye, C., I. Hurbain, D. Tenza, J.-B. Sibarita, S. Uzan-Gafsou, H. Ohno, W.J.C. Geerts, A.J. Verkleij, J. Salamero, M.S. Marks, and G. Raposo. 2009. AP-1 and KIF13A coordinate endosomal sorting and positioning during melanosome biogenesis. J. Cell Biol. 187:247–264. doi:10.1083/jcb.200907122.

Delevoye, C., S. Miserey-Lenkei, G. Montagnac, F. Gilles-Marsens, P. Paul-Gilloteaux, F. Giordano, F. Waharte, M.S. Marks, B. Goud, and G. Raposo. 2014. Recycling endosome tubule morphogenesis from sorting endosomes requires the kinesin motor KIF13A. Cell Rep. 6:445–454. doi:10.1016/j.celrep.2014.01.002.

Dell’Angelica, E.C., and J.S. Bonifacino. 2019. Coatopathies: Genetic Disorders of Protein Coats. Annu. Rev. Cell Dev. Biol. 35:1–38. doi:10.1146/annurev-cellbio-100818-125234.

Duncan, M.C. 2022. New directions for the clathrin adaptor AP-1 in cell biology and human disease. Curr. Opin. Cell Biol. 76:102079. doi:10.1016/j.ceb.2022.102079.

Eichel, K., T. Uenaka, V. Belapurkar, R. Lu, S. Cheng, J.S. Pak, C.A. Taylor, T.C. Südhof, R. Malenka, M. Wernig, E. Özkan, D. Perrais, and K. Shen. 2022. Endocytosis in the axon initial segment maintains neuronal polarity. Nature. 489:47–54. doi:10.1038/s41586-022-05074-5.

Encalada, S.E., L. Szpankowski, C. Xia, and L.S.B. Goldstein. 2011. Stable kinesin and dynein assemblies drive the axonal transport of mammalian prion protein vesicles. Cell. 144:551–565. doi:10.1016/j.cell.2011.01.021.

Encell, L.P. 2012. Development of a Dehalogenase-Based Protein Fusion Tag Capable of Rapid, Selective and Covalent Attachment to Customizable Ligands. Curr. Chem. Genomics. 6:55–71. doi:10.2174/1875397301206010055.

Etoh, K., and M. Fukuda. 2019. Rab10 regulates tubular endosome formation through KIF13A and KIF13B motors. J. Cell Sci. 132:jcs226977. doi:10.1242/jcs.226977.

Fang, C., H. Decker, and G. Banker. 2014. Axonal transport plays a crucial role in mediating the axon-protective effects of NmNAT. Neurobiol. Dis. 68:78–90. doi:10.1016/j.nbd.2014.04.013.

Farías, G.G., L. Cuitino, X. Guo, X. Ren, M. Jarnik, R. Mattera, and J.S. Bonifacino. 2012. Signal-mediated, AP-1/clathrin-dependent sorting of transmembrane receptors to the somatodendritic domain of hippocampal neurons. Neuron. 75:810–823. doi:10.1016/j.neuron.2012.07.007.

Farías, G.G., C.M. Guardia, D.J. Britt, X. Guo, and J.S. Bonifacino. 2015. Sorting of Dendritic and Axonal Vesicles at the Pre-axonal Exclusion Zone. Cell Rep. 13:1221–1232. doi:10.1016/j.celrep.2015.09.074.

Farías, G.G., C.M. Guardia, R. De Pace, D.J. Britt, and J.S. Bonifacino. 2017. BORC/kinesin-1 ensemble drives polarized transport of lysosomes into the axon. Proc. Natl. Acad. Sci. 201616363. doi:10.1073/pnas.1616363114.

Ford, C., A. Parchure, J. von Blume, and C.G. Burd. 2021. Cargo sorting at the trans-Golgi network at a glance. J. Cell Sci. 134:1–9. doi:10.1242/jcs.259110.

Frank, M., C.G. Citarella, G.B. Quinones, and M. Bentley. 2020. A Novel Labeling Strategy Reveals That Myosin Va and Myosin Vb Bind the Same Dendritically Polarized Vesicle Population. Traffic. 21:689–701. doi:10.1111/tra.12764.

Frank, M., A.T. Nabb, S.P. Gilbert, and M. Bentley. 2022. Propofol attenuates kinesin-mediated axonal vesicle transport and fusion. Mol. Biol. Cell. 33:ar119. doi:10.1091/mbc.E22-07-0276.

Fu, M.M., and E.L.F. Holzbaur. 2013. JIP1 regulates the directionality of APP axonal transport by coordinating kinesin and dynein motors. J. Cell Biol. 202:495–508. doi:10.1083/jcb.201302078.

Fukuda, Y., M.F. Pazyra-Murphy, E.S. Silagi, O.E. Tasdemir-Yilmaz, Y. Li, L. Rose, Z.C. Yeoh, N.E. Vangos, E.A. Geffken, H.S. Seo, G. Adelmant, G.H. Bird, L.D. Walensky, J.A. Marto, S. Dhe-Paganon, and R.A. Segal. 2021. Binding and transport of sfpq-rna granules by kif5a/klc1 motors promotes axon survival. J. Cell Biol. 220. doi:10.1083/jcb.202005051.

Futerman, A.H., and G.A. Banker. 1996. The economics of neurite outgrowth - The addition of new membrane to growing axons. Trends Neurosci. 19:144–149. doi:10.1016/S0166-2236(96)80025-7.

Ganguly, A., X. Han, U. Das, L. Wang, J. Loi, J. Sun, D. Gitler, G. Caillol, C. Leterrier, J.R. Yates, and S. Roy. 2017. Hsc70 chaperone activity is required for the cytosolic slow axonal transport of synapsin. J. Cell Biol. 216:2059–2074. doi:10.1083/jcb.201604028.

Ganguly, A., Y. Tang, L. Wang, K. Ladt, J. Loi, B. Dargent, C. Leterrier, and S. Roy. 2015. A dynamic formin-dependent deep F-actin network in axons. J. Cell Biol. 104:401–417. doi:10.1083/jcb.201506110.

Garbouchian, A., A. Montgomery, S.P. Gilbert, and M. Bentley. 2022. KAP is the neuronal organelle adaptor for Kinesin-2 KIF3AB and KIF3AC. Mol. Biol. Cell. 33. doi:10.1091/mbc.e22-08-0336.

Ghosh, P., and S. Kornfeld. 2003. AP-1 binding to sorting signals and release from clathrin-coated vesicles is regulated by phosphorylation. J. Cell Biol. 160:699–708. doi:10.1083/jcb.200211080.

Goodman, L., D. Baddeley, W. Ambroziak, C.L. Waites, C.C. Garner, C. Soeller, and J.M. Montgomery. 2017. N-terminal SAP97 isoforms differentially regulate synaptic structure and postsynaptic surface pools of AMPA receptors. Hippocampus. 27:668–682. doi:10.1002/hipo.22723.

Grimm, J.B., B.P. English, J. Chen, J.P. Slaughter, Z. Zhang, A. Revyakin, R. Patel, J.J. Macklin, D. Normanno, R.H. Singer, T. Lionnet, and L.D. Lavis. 2015. A general method to improve fluorophores for live-cell and single-molecule microscopy. Nat. Methods. 12:244–250. doi:10.1038/nmeth.3256.

Guardia, C.M., R. De Pace, R. Mattera, and J.S. Bonifacino. 2018. Neuronal functions of adaptor complexes involved in protein sorting. Curr. Opin. Neurobiol. 51:103–110. doi:10.1016/j.conb.2018.02.021.

Gutiérrez, Y., S. López-García, A. Lario, S. Gutiérrez-Eisman, C. Delevoye, and J.A. Esteban. 2021. Kif13a drives ampa receptor synaptic delivery for long-term potentiation via endosomal remodeling. J. Cell Biol. 220. doi:10.1083/jcb.202003183.

Hanada, T., L. Lin, E. V Tibaldi, E.L. Reinherz, and A.H. Chishti. 2000. GAKIN, a novel kinesin-like protein associates with the human homologue of the Drosophila discs large tumor suppressor in T lymphocytes. J. Biol. Chem. 275:28774–28784. doi:10.1074/jbc.M000715200.

Harding, C., J. Heuser, and P. Stahl. 1983. Receptor-mediated endocytosis of transferrin and recycling of the transferrin receptor in rat reticulocytes. J. Cell Biol. 97:329–39. doi:10.1083/jcb.97.2.329.

Hoo, L.S., C.D. Banna, C.M. Radeke, N. Sharma, M.E. Albertolle, S.H. Low, T. Weimbs, and C.A. Vandenberg. 2016. The SNARE Protein Syntaxin 3 Confers Specificity for Polarized Axonal Trafficking in Neurons. PLoS One. 11:e0163671. doi:10.1371/journal.pone.0163671.

Horiguchi, K., T. Hanada, Y. Fukui, and A.H. Chishti. 2006. Transport of PIP3 by GAKIN, a kinesin-3 family protein, regulates neuronal cell polarity. J. Cell Biol. 174:425–436. doi:10.1083/jcb.200604031.

Horikawa, H.P.M., M. Kneussel, O. El Far, and H. Betz. 2002. Interaction of synaptophysin with the AP-1 adaptor protein γ-adaptin. Mol. Cell. Neurosci. 21:454–462. doi:10.1006/mcne.2002.1191.

Horton, A.C., B. Rácz, E.E. Monson, A.L. Lin, R.J. Weinberg, and M.D. Ehlers. 2005. Polarized secretory trafficking directs cargo for asymmetric dendrite growth and morphogenesis. Neuron. 48:757–771. doi:10.1016/j.neuron.2005.11.005.

Huang, C.F., and G. Banker. 2012. The Translocation Selectivity of the Kinesins that Mediate Neuronal Organelle Transport. Traffic. 13:549–564. doi:10.1111/j.1600-0854.2011.01325.x.

Jacobson, C., B. Schnapp, and G.A. Banker. 2006. A change in the selective translocation of the Kinesin-1 motor domain marks the initial specification of the axon. Neuron. 49:797– 804. doi:10.1016/j.neuron.2006.02.005.

Jain, S., G.G. Farias, and J.S. Bonifacino. 2015. Polarized sorting of the copper transporter ATP7B in neurons mediated by recognition of a dileucine signal by AP-1. Mol. Biol. Cell. 26:218–228. doi:10.1091/mbc.E14-07-1177.

Janssen, A.F.J., R.P. Tas, P. van Bergeijk, R. Oost, C.C. Hoogenraad, and L.C. Kapitein. 2017. Myosin-V Induces Cargo Immobilization and Clustering at the Axon Initial Segment. Front. Cell. Neurosci. 11:260. doi:10.3389/fncel.2017.00260.

Janssen, A.F.J., P.S. Wulf, L.C. Kapitein, and C.C. Hoogenraad. 2016. Three-Step Model for Polarized Sorting of KIF17 into Dendrites. Curr Biol. 26:1705–1712. doi:10.1016/j.cub.2016.04.057.

Jareb, M., and G. Banker. 1997. Inhibition of axonal growth by brefeldin A in hippocampal neurons in culture. J. Neurosci. 17:8955–8963. doi:10.1523/jneurosci.17-23-08955.1997.

Jenkins, B., H. Decker, M. Bentley, J. Luisi, and G. Banker. 2012. A novel split kinesin assay identifies motor proteins that interact with distinct vesicle populations. J. Cell Biol. 198:749– 761. doi:10.1083/jcb.201205070.

Jensen, C.S., S. Watanabe, H.B. Rasmussen, N. Schmitt, S.-P. Olesen, N.A. Frost, T.A. Blanpied, and H. Misonou. 2014. Specific sorting and post-Golgi trafficking of dendritic potassium channels in living neurons. J. Biol. Chem. 289:10566–10581. doi:10.1074/jbc.M113.534495.

de Jong, G., J.P. van Dijk, and H.G. van Eijk. 1990. The biology of transferrin. Clin. Chim. Acta. 190:1–46. doi:10.1016/0009-8981(90)90278-z.

Kaech, S., and G. Banker. 2006. Culturing hippocampal neurons. Nat. Protoc. 1:2406–2415. doi:10.1038/nprot.2006.356.

Kaech, S., C.F. Huang, and G. Banker. 2012a. Short-term high-resolution imaging of developing hippocampal neurons in culture. Cold Spring Harb. Protoc. 7:340–343. doi:10.1101/pdb.prot068247.

Kaech, S., C.F. Huang, and G. Banker. 2012b. General considerations for live imaging of developing hippocampal neurons in culture. Cold Spring Harb. Protoc. 7:312–318. doi:10.1101/pdb.ip068221.

Kamiguchi, H., K.E. Long, M. Pendergast, A.W. Schaefer, I. Rapoport, T. Kirchhausen, and V. Lemmon. 1998. The neural cell adhesion molecule L1 interacts with the AP-2 adaptor and is endocytosed via the clathrin-mediated pathway. J. Neurosci. 18:5311–5321. doi:10.1523/jneurosci.18-14-05311.1998.

Kapitein, L.C., M.A. Schlager, M. Kuijpers, P.S. Wulf, M. van Spronsen, F.C. MacKintosh, and C.C. Hoogenraad. 2010. Mixed Microtubules Steer Dynein-Driven Cargo Transport into Dendrites. Curr. Biol. 20:290–299. doi:10.1016/j.cub.2009.12.052.

Kelliher, M.T., Y. Yue, A. Ng, D. Kamiyama, B. Huang, K.J. Verhey, and J. Wildonger. 2018. Autoinhibition of kinesin-1 is essential to the dendrite-specific localization of Golgi outposts. J. Cell Biol. 217:2531–2547. doi:10.1083/jcb.201708096.

Keren-Kaplan, T., and J.S. Bonifacino. 2021. ARL8 Relieves SKIP Autoinhibition to Enable Coupling of Lysosomes to Kinesin-1. Curr. Biol. 31:540–554.e5. doi:10.1016/j.cub.2020.10.071.

Kuijpers, M., D. van de Willige, A. Freal, A. Chazeau, M.A. Franker, J. Hofenk, R.J.C. Rodrigues, L.C. Kapitein, A. Akhmanova, D. Jaarsma, and C.C. Hoogenraad. 2016. Dynein Regulator NDEL1 Controls Polarized Cargo Transport at the Axon Initial Segment. Neuron. 89:461–471. doi:10.1016/j.neuron.2016.01.022.

Leterrier, C., and B. Dargent. 2014. No Pasaran! Role of the axon initial segment in the regulation of protein transport and the maintenance of axonal identity. Semin. Cell Dev. Biol. 27:44–51. doi:10.1016/j.semcdb.2013.11.001.

Lewis, T.L., T. Mao, and D.B. Arnold. 2011. A role for Myosin VI in the localization of axonal proteins. PLoS Biol. 9. doi:10.1371/journal.pbio.1001021.

Lewis, T.L., T. Mao, K. Svoboda, and D.B. Arnold. 2009. Myosin-dependent targeting of transmembrane proteins to neuronal dendrites. Nat. Neurosci. 12:568–576. doi:10.1038/nn.2318.

Li, P., S.A. Merrill, E.M. Jorgensen, and K. Shen. 2016. Two Clathrin Adaptor Protein Complexes Instruct Axon-Dendrite Polarity. Neuron. 90:564–580. doi:10.1016/j.neuron.2016.04.020.

Lo, K.Y., A. Kuzmin, S.M. Unger, J.D. Petersen, and M.A. Silverman. 2011. KIF1A is the primary anterograde motor protein required for the axonal transport of dense-core vesicles in cultured hippocampal neurons. Neurosci. Lett. 491:168–173. doi:10.1016/j.neulet.2011.01.018.

Los, G. V, L.P. Encell, M.G. McDougall, D.D. Hartzell, N. Karassina, C. Zimprich, M.G. Wood, R. Learish, R.F. Ohana, M. Urh, D. Simpson, J. Mendez, K. Zimmerman, P. Otto, G. Vidugiris, J. Zhu, A. Darzins, D.H. Klaubert, R.F. Bulleit, and K. V Wood. 2008. HaloTag: a novel protein labeling technology for cell imaging and protein analysis. ACS Chem. Biol. 3:373–82. doi:10.1021/cb800025k.

Maritzen, T., and V. Haucke. 2010. Gadkin: A novel link between endosomal vesicles and microtubule tracks. Commun. Integr. Biol. 3:299–302. doi:10.4161/cib.3.4.11835.

El Meskini, R., L. Jin, R. Marx, A. Bruzzaniti, J. Lee, R.B. Emeson, and R.E. Mains. 2001. A signal sequence is sufficient for green fluorescent protein to be routed to regulated secretory granules. Endocrinology. 142:864–873. doi:10.1210/endo.142.2.7929.

Mills, J., T. Hanada, Y. Hase, L. Liscum, and A.H. Chishti. 2019. LDL receptor related protein 1 requires the I3 domain of discs-large homolog 1/DLG1 for interaction with the kinesin motor protein KIF13B. Biochim. Biophys. Acta - Mol. Cell Res. 1866:118552. doi:10.1016/j.bbamcr.2019.118552.

Montgomery, A., A. Garbouchian, and M. Bentley. 2022. Visualizing Vesicle-Bound Kinesins in Cultured Hippocampal Neurons. Methods Mol. Biol. 2431:239–247. doi:10.1007/978-1-0716-1990-2_12.

Moore, F.B., and J.D. Baleja. 2012. Molecular remodeling mechanisms of the neural somatodendritic compartment. Biochim. Biophys. Acta. 1823:1720–30. doi:10.1016/j.bbamcr.2012.06.006.

Nabb, A.T., and M. Bentley. 2022. NgCAM and VAMP2 reveal that direct delivery and dendritic degradation maintain axonal polarity. Mol. Biol. Cell. 33:ar3. doi:10.1091/mbc.E21-08-0425.

Nabb, A.T., M. Frank, and M. Bentley. 2020. Smart motors and cargo steering drive kinesin-mediated selective transport. Mol. Cell. Neurosci. 103:103464. doi:10.1016/j.mcn.2019.103464.

Nakagawa, T., M. Setou, D.H. Seog, K. Ogasawara, N. Dohmae, K. Takio, and N. Hirokawa. 2000. A novel motor, KIF13A, transports mannose-6-phosphate receptor to plasma membrane through direct interaction with AP-1 complex. Cell. 103:569–581. doi:10.1016/S0092-8674(00)00161-6.

Nakata, T., and N. Hirokawa. 2003. Microtubules provide directional cues for polarized axonal transport through interaction with kinesin motor head. J. Cell Biol. 162:1045–1055. doi:10.1083/jcb.200302175.

Nakata, T., S. Niwa, Y. Okada, F. Perez, and N. Hirokawa. 2011. Preferential binding of a kinesin-1 motor to GTP-tubulin-rich microtubules underlies polarized vesicle transport. J. Cell Biol. 194:245–255. doi:10.1083/jcb.201104034.

Pearse, B.M.F., and M.S. Robinson. 1990. Clathrin, adaptors, and sorting. Annu. Rev. Cell Biol.

Petersen, J.D., S. Kaech, and G. Banker. 2014. Selective microtubule-based transport of dendritic membrane proteins arises in concert with axon specification. J. Neurosci. 34:4135–47. doi:10.1523/jneurosci.3779-13.2014.

Radler, M.R., A. Suber, and E.T. Spiliotis. 2020. Spatial control of membrane traffic in neuronal dendrites. Mol. Cell. Neurosci. 105:103492. doi:10.1016/j.mcn.2020.103492.

Ramazanov, B.R., M.L. Tran, and J. von Blume. 2021. Sending out molecules from the TGN. Curr. Opin. Cell Biol. 71:55–62. doi:10.1016/j.ceb.2021.02.005.

Ren, X., G.G. Farías, B.J. Canagarajah, J.S. Bonifacino, and J.H. Hurley. 2013. Structural Basis for Recruitment and Activation of the AP-1 Clathrin Adaptor Complex by Arf1. Cell. 152:755–767. doi:10.1016/j.cell.2012.12.042.

Robinson, M.S., D.A. Sahlender, and S.D. Foster. 2010. Rapid inactivation of proteins by rapamycin-induced rerouting to mitochondria. Dev. Cell. 18:324–331. doi:10.1016/j.devcel.2009.12.015.

Rumbaugh, G., G.M. Sia, C.C. Garner, and R.L. Huganir. 2003. Synapse-associated protein-97 isoform-specific regulation of surface AMPA receptors and synaptic function in cultured neurons. J. Neurosci. 23:4567–4576. doi:10.1523/jneurosci.23-11-04567.2003.

Sampo, B., S. Kaech, S. Kunz, and G. Banker. 2003. Two distinct mechanisms target membrane proteins to the axonal surface. Neuron. 37:611–24. doi:10.1016/s0896-6273(03)00058-8.

Sanger, A., J. Hirst, A.K. Davies, and M.S. Robinson. 2019. Adaptor protein complexes and disease at a glance. J. Cell Sci. 132. doi:10.1242/jcs.222992.

Sans, N., C. Racca, R.S. Petralia, Y.X. Wang, J. McCallum, and R.J. Wenthold. 2001. Synapse-associated protein 97 selectively associates with a subset of AMPA receptors early in their biosynthetic pathway. J. Neurosci. 21:7506–7516. doi:10.1523/jneurosci.21-19-07506.2001.

Schaefer, A.W., Y. Kamei, H. Kamiguchi, E. V. Wong, I. Rapoport, T. Kirchhausen, C.M. Beach, G. Landreth, S.K. Lemmon, and V. Lemmon. 2002. L1 endocytosis is controlled by a phosphorylation-dephosphorylation cycle stimulated by outside-in signaling by L1. J. Cell Biol. 157:1223–1232. doi:10.1083/jcb.200203024.

Schmidt, M.R., T. Maritzen, V. Kukhtina, V.A. Higman, L. Doglio, N.N. Barak, H. Strauss, H. Oschkinat, C.G. Dotti, and V. Haucke. 2009. Regulation of endosomal membrane traffic by a Gadkin/AP-1/kinesin KIF5 complex. Proc. Natl. Acad. Sci. U. S. A. 106:15344–9. doi:10.1073/pnas.0904268106.

Shih, W., A. Gallusser, and T. Kirchhausen. 1995. A clathrin-binding site in the hinge of the β2 chain of mammalian AP-2 complexes. J. Biol. Chem. 270:31083–31090. doi:10.1074/jbc.270.52.31083.

Silverman, M.A., S. Kaech, M. Jareb, M.A. Burack, L. Vogt, P. Sonderegger, and G. Banker. 2001. Sorting and directed transport of membrane proteins during development of hippocampal neurons in culture. Proc. Natl. Acad. Sci. U. S. A. 98:7051–7057. doi:10.1073/pnas.111146198.

Tong, Y., W. Tempel, H. Wang, K. Yamada, L. Shen, G.A. Senisterra, F. MacKenzie, A.H. Chishti, and H.W. Park. 2010. Phosphorylation-independent dual-site binding of the FHA domain of KIF13 mediates phosphoinositide transport via centaurin α1. Proc. Natl. Acad. Sci. U. S. A. 107:20346–20351. doi:10.1073/pnas.1009008107.

Vale, R.D., T.S. Reese, and M.P. Sheetz. 1985a. Identification of a novel force-generating protein, kinesin, involved in microtubule-based motility. Cell. 42:39–50. doi:10.1016/s0092-8674(85)80099-4.

Vale, R.D., B.J. Schnapp, T. Mitchison, E. Steuer, T.S. Reese, and M.P. Sheetz. 1985b. Different axoplasmic proteins generate movement in opposite directions along microtubules in vitro. Cell. 43:623–32. doi:10.1016/0092-8674(85)90234-x.

Venkateswarlu, K., T. Hanada, and A.H. Chishti. 2005. Centaurin-α1 interacts directly with kinesin motor protein KIF13B. J. Cell Sci. 118:2471–2484. doi:10.1242/jcs.02369.

Waites, C.L., C.G. Specht, K. Härtel, S. Leal-Ortiz, D. Genoux, D. Li, R.C. Drisdel, O. Jeyifous, J.E. Cheyne, W.N. Green, J.M. Montgomery, and C.C. Garner. 2009. Synaptic SAP97 isoforms regulate AMPA receptor dynamics and access to presynaptic glutamate. J. Neurosci. 29:4332–45. doi:10.1523/jneurosci.4431-08.2009.

Wang, X., and T.L. Schwarz. 2009. The mechanism of Ca2+ - dependent regulation of kinesin-mediated mitochondrial motility. Cell. 136:163–74. doi:10.1016/j.cell.2008.11.046.

Wang, Z., J.G. Edwards, N. Riley, D.W. Provance, R. Karcher, X. Li, I.G. Davison, M. Ikebe, J.A. Mercer, J.A. Kauer, and M.D. Ehlers. 2008. Myosin Vb Mobilizes Recycling Endosomes and AMPA Receptors for Postsynaptic Plasticity. Cell. 135:535–548. doi:10.1016/j.cell.2008.09.057.

Watanabe, K., S. Al-Bassam, Y. Miyazaki, T.J. Wandless, P. Webster, and D.B. Arnold. 2012. Networks of polarized actin filaments in the axon initial segment provide a mechanism for sorting axonal and dendritic proteins. Cell Rep. 2:1546–1553. doi:10.1016/j.celrep.2012.11.015.

Wilde, A., and F.M. Brodsky. 1996. In vivo phosphorylation of adaptors regulates their interaction with clathrin. J. Cell Biol. 135:635–45. doi:10.1083/jcb.135.3.635.

Wisco, D., E.D. Anderson, M.C. Chang, C. Norden, T. Boiko, H. Fölsch, and B. Winckler. 2003. Uncovering multiple axonal targeting pathways in hippocampal neurons. J. Cell Biol. 162:1317–1328. doi:10.1083/jcb.200307069.

Yamada, K.H., T. Hanada, and A.H. Chishti. 2007. The effector domain of human Dlg tumor suppressor acts as a switch that relieves autoinhibition of kinesin-3 motor GAKIN/KIF13B. Biochemistry. 46:10039–45. doi:10.1021/bi701169w.

Yang, R., Z. Bostick, A. Garbouchian, J. Luisi, G. Banker, and M. Bentley. 2019. A novel strategy to visualize vesicle-bound kinesins reveals the diversity of kinesin-mediated transport. Traffic. 20:851–866. doi:10.1111/tra.12692.

Yap, C.C., R.L. Nokes, D. Wisco, E. Anderson, H. Folsch, and B. Winckler. 2008. Pathway selection to the axon depends on multiple targeting signals in NgCAM. J Cell Sci. 121:1514–1525. doi:10.1242/jcs.022442.

Yoon, Y.J., B. Wu, A.R. Buxbaum, S. Das, A. Tsai, B.P. English, J.B. Grimm, L.D. Lavis, and R.H. Singer. 2016. Glutamate-induced RNA localization and translation in neurons. Proc. Natl. Acad. Sci. U. S. A. 113:E6877–E6886. doi:10.1073/pnas.1614267113.

Yoshimura, Y., T. Terabayashi, and H. Miki. 2010. Par1b/MARK2 phosphorylates kinesin-like motor protein GAKIN/KIF13B to regulate axon formation. Mol. Cell. Biol. 30:2206–19. doi:10.1128/MCB.01181-09.

Yu, H.L., Y. Peng, Y. Zhao, Y.S. Lan, B. Wang, L. Zhao, D. Sun, J.X. Pan, Z.Q. Dong, L. Mei, Y.Q. Ding, X.J. Zhu, and W.C. Xiong. 2020. Myosin X interaction with KIF13B, a crucial pathway for Netrin-1-Induced axonal development. J. Neurosci. 40:9169–9185. doi:10.1523/JNEUROSCI.0929-20.2020.

Zheng, Z., and J. Keifer. 2014. Sequential delivery of synaptic GluA1- and GluA4-containing AMPa receptors (AMPARs) by SAP97 anchored protein complexes in classical conditioning. J. Biol. Chem. 289:10540–10550. doi:10.1074/jbc.M113.535179.

Zhou, R., S. Niwa, L. Guillaud, Y. Tong, and N. Hirokawa. 2013. A Molecular Motor, KIF13A, Controls Anxiety by Transporting the Serotonin Type 1A Receptor. Cell Rep. 3:509–519. doi:10.1016/j.celrep.2013.01.014.

